# The first horse gut microbiome gene catalog reveals that rare microbiome ensures better cardiovascular fitness in endurance horses

**DOI:** 10.1101/2022.01.24.477461

**Authors:** Núria Mach, Cédric Midoux, Sébastien Leclercq, Samuel Pennarun, Laurence Le Moyec, Olivier Rué, Céline Robert, Guillaume Sallé, Eric Barrey

**Author notes:** These authors contributed equally (Guillaume Sallé, Eric Barrey). **Materials and correspondence:** Núria Mach.

## Abstract

Emerging evidence indicates that the gut microbiome contributes to endurance exercise performance, but the extent of their functional and metabolic potential remains unknown. Using elite endurance horses as a model system for exercise responsiveness, we built the first equine gut microbial gene catalog comprising more than 25 million non-redundant genes representing 4,696 genera spanning 95 phyla. The unprecedented resolution unrevealed functional pathways relevant for both the structure of the microbiome and the host health and recovered 369 novel metagenome-assembled bacterial genomes, providing useful reference for future studies. Integration of microbial and host omic datasets suggested that microbiomes harboring rare species were functionally dissimilar from those enriched in *Lachnospiraceae* taxa. Moreover, they offered expanded metabolic pathways to fine-tune the cardiovascular capacity through mitochondria-mediated mechanisms. The results identify an associative link between horse endurance capability and its microbiome gene function, laying the basis for nutritional interventions that could benefit endurance athletes.

## INTRODUCTION

Endurance athletes undergo prolonged cardiovascular exercise and withstand physiological stress that disrupts the body’s homeostasis. This, in turn, overwhelms organs and the system’s normal function ^1,2^. The ability to run for long distances at high speed is scarcely distributed across land mammals. Through years of selective breeding, Arabian horses have gained built-in biological mechanisms to run at more than 160 km at 20 km/h, an effort comparable to that of marathon or ultra-marathon runners ^3^. The effective body heat dissipation and the ability to endure extreme exercise enable this breed to have outstanding endurance capabilities ^4^.

Endurance exercise performance is primarily limited by cardiovascular fitness, exercise economy and the ability to sustain work without either excessive blood lactate levels or fatigue ^5^. The athletes with the greatest improved cardiovascular fitness and fatigue resistance often succeed in competitions ^5^.

Undoubtedly, endurance exercise performance entails complex multifactorial processes whose mechanisms are still not fully understood. New evidence has shown that the gut microbiome and its associated metabolites impact host athletic performance during endurance racing in humans ^6,7^. The gut microbiome produces thousands of unique small molecules that can potentially affect many aspects of physiology such as regulating immunity, hydration and redox reactions as well as shaping the gut-brain axis that affects fatigue and stress perception ^2,8–12^.

These metabolites can act locally in the intestine or can accumulate up to millimolar concentrations in different body fluids ^13^. Higher microbial diversity has been correlated with improved cardiorespiratory fitness and performance in marathon runners regardless of the sex, age, body mass index and diet ^14^. Other reports have also found significant associations between the cardiovascular capacity, as assessed by the maximum oxygen consumption and the Firmicutes-Bacteroidetes ratio ^15^ or reduced levels of fecal *Eubacterium* spp. ^16^. Deeper characterization of the links between athletic performance and the gut microbiome revealed that the single bacterium *Veillonella atypica* is required to enhance athletic performance in treadmill mice experiments ^6,7^.

While current knowledge of the relationships between gut microbiome and endurance performance are in their infancy in humans, controlling for known confounding factors (such as diet, training loads, medications, occurring illnesses, environment and genetic background) has proven difficult. In this respect, Arabian horses emerge as a suitable *in vivo* model for characterizing the microbiome adaptations to endurance exercise due to their natural aptitude for athletic performance, the homogeneity of their genetic and environmental backgrounds and the relative ease of sampling during endurance races. Furthermore, recent findings suggest that gut microbial metabolites in endurance horses act as mitochondria function regulators that prevent hypoglycemia ^17^, which is the limiting factor for fatigue onset and thus, performance.

Despite these findings, if and how gut microbiome functions are responsible for better adaptations to fatigue resistance, as well as success in athletic performances are not well understood. To address this gap, we have built the first gene catalog of the equine gut microbiome in elite endurance horses. Our results expanded the current representation of the equine gut microbiome with more than 25 millions of non-redundant genes identified and 369 new metagenome-assembled genomes (MAGs). Moreover, by using the holo-omic approach that incorporates multi-omic data from host and microbiome domains we have shown that the gut microbiome composition and functions and the mitochondria activity are key determinants for cardiovascular fitness. Rare microbes and their pool of genetic resources likely offered metabolic pathways that fine-tuned mitochondrial function and consequently confer enhanced cardiovascular capacity compared to microbial ecosystems with reduced diversity but higher abundance of the *Lachnospiraceae* family.

## RESULTS

### Building the first horse gut microbiome gene catalog

We constructed a microbial gene catalogue from the feces of 11 highly trained endurance horses (Suppl Table S1). After quality filtering and host sequences decontamination, 1,124 millions of high-quality clean paired reads were available, with an average sequencing depth per sample similar to that used for the construction of chicken ^18^ and bovine ^19^ gut gene catalog (*n* = of 102 millions of paired reads per sample on average; Suppl Table S2). These data were *de novo* assembled (total assembly size of 21.68 Gb; Suppl Fig S1a) to build a non-redundant gene catalog of 25,250,066 genes with an average length of 618 bp (Suppl Table S2 and Fig S1a-f). Individual horses harbored around half of these genes (*n* = 11,809,713 genes) on average (Fig 1a).

**Figure 1.**
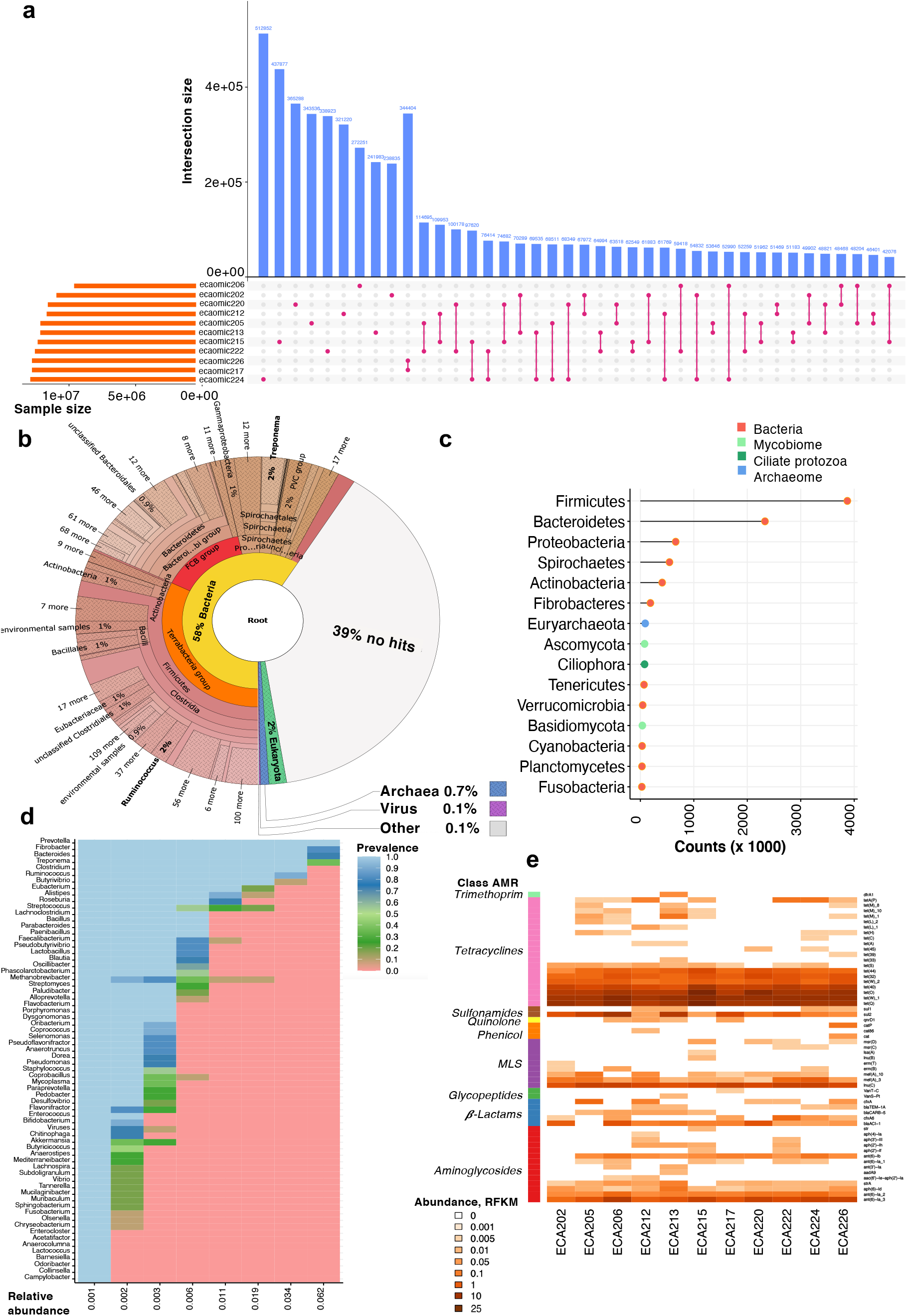
Description of the first horse gut gene catalog: core microbiome and taxonomic annotation. (a) Contribution of different sample sources to gene content of the horse gut catalog. Vertical purple and blue bars represent the number of genes present in only one sample or shared between pairs of samples, respectively. Horizontal orange bars in the lower panel indicate the total number of genes contained in each sample; (b)Visualization of the taxonomic assignment of Illumina reads in a Krona plot using the software tool Kaiju; (c) Lollipop plot showing the read counts identified by the Kaiju resolved at the phylum level. Dots are colored by kingdom; (d) Heatmap depicting the core phylome and their prevalence at different detection thresholds (relative abundance). The percentage of shared items and the proportion of shared samples are represented on the *y*- and *x*-axis, respectively; (e) Heatmap showing the normalized counts of antimicrobial resistance (AMR) genes for each individual based on ResFinder database

Taxonomic assignments were made from clean read pairs searching for maximum exact matches between translated sequences and a reference database ^20^. This approach yielded an annotation for 61% of all sequences (Fig 1b) and revealed a diverse community of 95 phyla encompassing 1,110 families and 4,696 genera. Bacteria (95%) defined most of the assemblage in terms of abundance and diversity, followed by a handful of eukaryotes (3.27%), archaea (1.14%) and viruses (0.15%). At the phylum level, the bacterial phyla Firmicutes (44.3%) and Bacteroidetes (26.5%) greatly outnumbered Proteobacteria (7.5%), Spirochaetes (6.2%), Actinobacteria (4.6%), Euryarchaeota (1.1%), Ciliophora (0.90%) and Ascomycota (0.90%; Fig 1c). Consistent with a recent metagenomic study in the wild Przewalski’s horses ^21^, Ascomycota and Basidiomycota were among the top Eukaryota phylum in the gut.

We then identified the most dominant microbial phylotypes. Dominant phylotypes (top 25% most common phylotypes sorted by their abundance and found in more than half of the samples) accounted for ∼94% of the total annotated sequences on average and were represented by 1,146 unique genera (Suppl Table S3). The great majority of these dominant microbes (85%) closely matched the most abundant bacterial genera recovered using 16S rRNA based prediction (relative abundance > 0.1%; Suppl Table S4 and Fig S1g). The core microbiome (Fig 1d) was defined by a narrow section of the phylomes (less than 0.15% of the overall microbial genera) that were highly abundant (38 to 54% of the sequences). Along with *Bacteroides* (6.66% ± 0.57) and *Prevotella* (9.41% ± 1.54), which have been found to be dominant genera in the gut of marathon and triathlon athletes, respectively ^22^, this core microbiome included species from *Fibrobacter* (9.44% ± 3.71), *Treponema* (5.96% ± 1.59), *Clostridium* (4.91% ± 0.49) and *Ruminococcus* (4.04% ± 0.79). All of them are in full agreement with the core microbiome of endurance horses inferred from 16S rRNA-based sequencing ^23–26^.

To gain functional insights, genes and functional modules were annotated using the non-supervised orthologous groups (EggNOG) database. We identified a total of 12,060 KEGG orthologous groups (KOs) and 137 different carbohydrate-active enzymes (CAZymes), which encompassed 44% (*n* = 11,132,404) and 3.38% (*n* = 665,235) of the gene catalog, respectively. The majority of KOs had functions related to essential microbial gut functions, including genetic information processing and signaling, carbohydrate and amino acid metabolism. However, we also identified a number of pathways (*i*.*e*., drug resistance, biosynthesis of secondary metabolites, endocrine system and neurodegenerative diseases) relevant for both the structure of the microbiome and the host health (Suppl Fig S2a). Most of the CAZYmes (85.4%; *n* = 568,462 genes) pertained to glycoside hydrolase (GH) and polysaccharide lyases (PL) families, highlighting the indispensable role of gut metagenome in complex dietary glycans metabolism (Suppl Fig S2b and Table S5).

Taking advantage of this deep shot-gun sequencing of the microbiome, we investigated the presence of antimicrobial resistance (AMR) genes. Considered horses harbored 57 clusters of AMR genes representing the major antibiotic classes, including tetracycline (*n* = 20), aminoglycosides (*n* = 20) and macrolides, lincosamides and streptogramins (MLS, *n* = 9; Fig 1e; Suppl Table S6). The overall AMR gene composition was similar to those of human ^27^ and livestock species such as cattle, pig and chicken, with a high abundance and prevalence of the Firmicutes and Bacteroidetes-associated tetracyclines resistance genes *tet*(W), *tet*(Q), *tet*(O), *tet*(40) and MLS resistance genes *lnu*(C) and *mef*(A) ^28–30^. Of note, we detected the extended spectrum of β-lactamase (ESBL) *bla*_ACI-1_ in 10 of the 11 horses. This AMR gene, found in several Negativicutes (Gram negative Firmicutes), is only rarely detected in animal or human gut microbiomes ^31^. Such unusual prevalence is likely linked to the presence of two Negativicutes genera, *Phascolarctobacterium* and *Selenomonas*, in the core microbiome of the studied horses (Fig 1d).

Lastly, a set of 372 non-redundant prokaryotic MAGs were constructed from the metagenomic sequencing at a threshold of > 50% completeness and contamination ≤ 10% (Suppl Table S7). Among these, 121 MAGs were estimated to be near complete; MAGs in this subset had minimal contamination (≤ 5%), high completeness (> 95%; Suppl Table S7). This MAG repertoire was assigned to 361 bacteria and 11 archaea, involving bacteria from the Bacteroidetes and Firmicutes phyla, followed by Spirochaetes, Euryarchaeota, Verrucomicrobia, Fibrobacteres and Cyanobacteria phylum (Suppl Fig S3ab). The abundance of genomes pertaining to Cyanobacteria, Proteobacteria and Verrucomicrobia phylum showed high rates of divergence between hosts (Suppl Fig S3c). This trend was bolstered at the lower taxonomic level, except for MAGs assigned to *Fibrobacter* spp. (Suppl Fig S3d). Of note, most MAGs (*n* = 369) have never been described before in horses to date ^32,33^, increasing the mappability of metagenomes and expanding our understanding of the horse microbiomes.

Altogether, the building of the gut gene catalog and MAGs repertoire expands current understanding of the equine gut microbiome. Its complexity and the abundance of genes associated with complex carbohydrate fermentation underscores the adaptation to a terrestrial herbivorous lifestyle while reminding the pervasive presence of AMR genes. Additionally, the identification of metagenome functional capacity primed for host health likely reflected the significant energy demands and tissue adaptations that occur during endurance exercise.

### Basal gut metagenome composition discriminates cardiovascular fitness

Building on the dominant microbial phylotypes, we investigated whether diversity, compositional and functional differences could classify samples according to athletic performance. First, the ordination of individual horse metagenomes using a non-metric multidimensional scaling of the dominant phylotypes identified two distinct groups of samples that recapitulated variation along the first axis (Fig 2a). A similar pattern was obtained with a principal coordinate analysis (PCoA, Suppl Fig S4a) and the two groups were also supported by permutational analysis of variance (PerMANOVA; *p* = 0.01, R^2^ = 0.3715). Cluster 1 (*n* = 3 horses) was represented by multiple taxa, involving ciliophora, methanogenic archaea as well as rare bacterial species from the Proteobacteria and Verrucomicrobia phylum (Fig 2b-c) and exhibited higher *α*-diversity (Shannon and inverse Simpson indices; *p* = 0.0134, Mann-Whitney *U* test, Fig 2d-e) despite the small sample size considered. In contrast to the overwhelming diversity observed in cluster 1, a skewed species abundance distribution, with predominance of phylotypes from the Firmicutes phylum (mainly *Lachnospiraceae* taxa) and *Fibrobacter* and *Treponema* genera defined cluster 2. Similarly, the sample distribution based on KOs and CAZymes profiles echoed that of the dominant phylotypes composition (Suppl Fig S4b-c, respectively).

**Figure 2.**
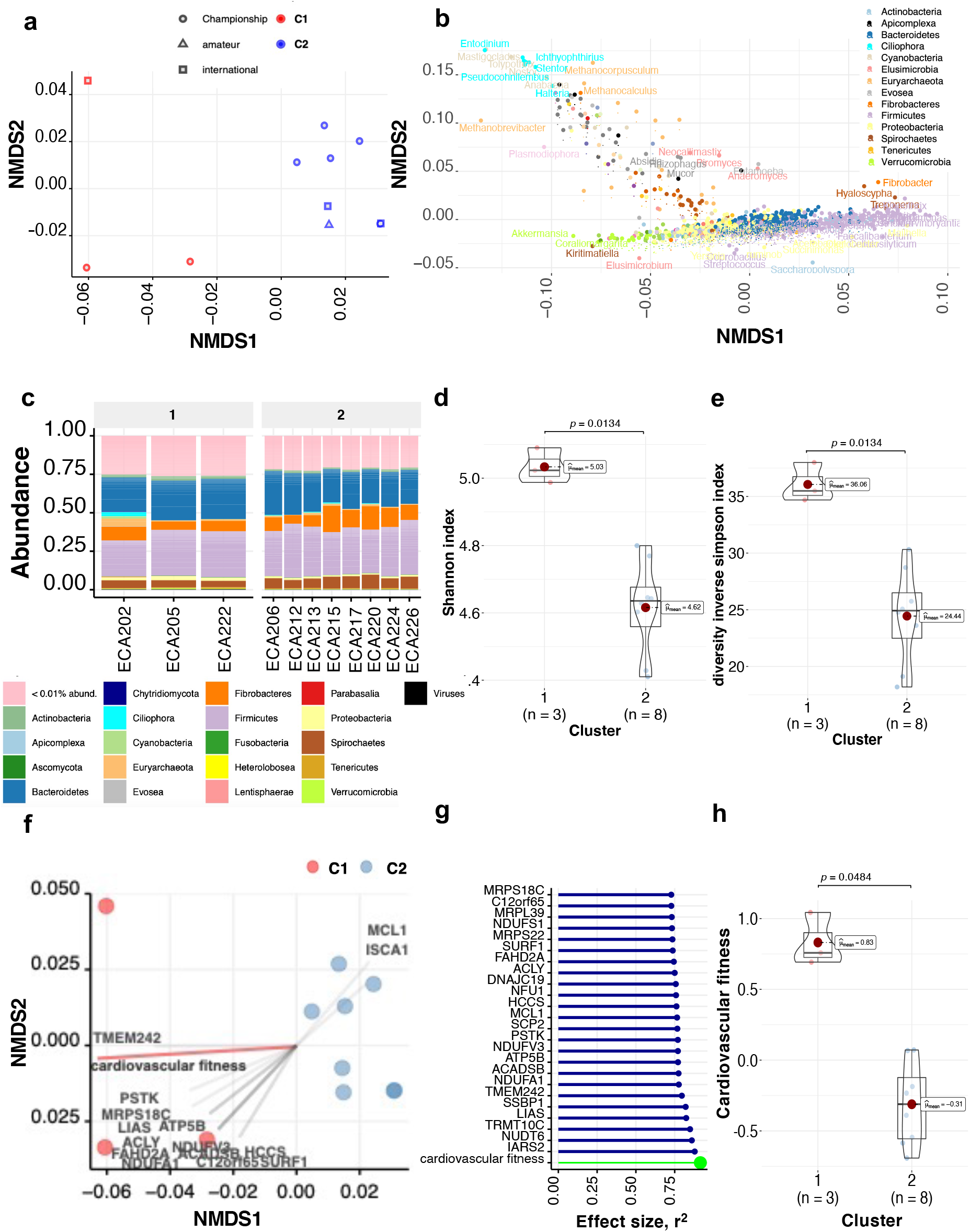
Gut microbiome composition and structure in endurance horses. (a) NMDS ordination analysis (Bray-Curtis distance) of dominant phylotypes composition. Points denote individual samples which are colored according to the clustering group. The shape of the dots indicates the competition level of horses; (b) Biplot values of the dominant phylotypes driving the NMDS ordination. The phylotypes contributing to the distinction between groups on at least one axis are depicted. Points are colored by phylum; (c) Taxonomic distribution of the relative abundance of phyla in each individual. Individuals are split by cluster; (d-e) Violin plot representing Shannon Diversity Index and inverse Simpson index, respectively. In all cases, colors indicate community classification, the community type 1 (red color) and community type 2 (blue color). Boxplots show median, 25^th^ and 75^th^ percentile, the whiskers indicate the minima and maxima and the points lying outside the whiskers of boxplots represent the outliers. Adjusted *p* values from Wilcoxon rank-sum tests; (f) NMDS ordination plot showing the covariates that contribute significantly to the variation of dominant phylotypes determined by “*envfit*” function. The arrows for each variable show the direction of the effect and are scaled by the unconditioned *r*^2^ value. Dots represent samples, which are colored according to the type of community: the community type 1 (red color) and community type 2 (blue color); (g) Effect sizes of the main variables affecting the NMDS ordination. The length of the horizontal bars shows the amount of variance (*r*^2^) explained by each covariate in the model. Covariates are colored according to the type of dataset: athletic performance are in green and mitochondrial related genes in blue; (h) Violin plot representing the cardiovascular fitness, which was calculated as a composite of post-exercise heart rate, cardiac recovery time and average speed during the race. Colors indicate community classification, the community type 1 (red color) and community type 2 (blue color) and boxplots show median, 25^th^ and 75^th^ percentile, the whiskers indicate the minima and maxima and the points lying outside the whiskers of boxplots represent the outliers. Adjusted *p* values from Wilcoxon rank-sum tests.

The macronutrient intake was not statistically different between horses from the two clusters (*p* > 0.05; Suppl Table S1). Therefore, we next tested the extent to which any horse physiological, metabolic or performance indicators best captured the distributions of microbial taxa. Host-centered omic and phenomics data, including transcriptomics, metabolomics, acylcarnitines and blood biochemical assay profiles as well as cardiovascular fitness parameters were used (Suppl Tables S8-S11, respectively). The cardiovascular fitness, a composite of post-exercise heart rate, cardiac recovery time and average speed during the race, was the principal contributor to the metagenome heterogeneity (*envfit*, R^2^ = 0.9192, adjusted *p* = 0.005; Suppl Table S12), outperforming the expression of several mitochondrial genes (Fig 2f-g). This composite parameter aggregated 39.64% of fecal microbiome community variation. Horses from cluster 1 had significantly higher cardiovascular fitness relative to that of cluster 2 members (*p* = 0.0484, Wilcoxon rank-sum test, Fig 2h) without incurring dramatic increases in blood lactate concentration (*p* = 0.9212, Wilcoxon rank-sum test), a proxy for glycolytic stress and disturbance in cellular homeostasis ^1^. The results hence indicated that - under the same prevailing environmental conditions and nutrient availability - individuals harboring cluster 1 type communities achieved improved cardiovascular capacity, that is lower post-exercise heart rates and faster cardiac recovery time at the veterinary inspection compared to individuals with cluster 2 type microbial communities. The cluster 2 individuals were harboring a less diverse gut microbiota with only a few rare species involved.

### An independent validation of findings confirms that *Lachnospiraceae* bacteria was associated with cardiovascular fitness and in highly trained equine athletes

To further confirm the association between the cardiovascular fitness and the gut microbiome composition found in the 11 elite horses based on their metagenome data, we analyzed the 16S rRNA sequence data from the gut microbiota of 22 independent highly trained endurance horses (Suppl Table S13 and S14). As with the study cohort, the microorganisms’ community profiles could be distinguished based on the horse’s cardiovascular fitness (adjusted *p* = 0.05; pairwise comparisons using PerMANOVA on a Bray-Curtis distance matrix, Suppl Fig S5a-c). Individuals with lower cardiovascular fitness harbored a few players belonging to Firmicutes phylum with higher abundance (adjusted *p* = 0.024, Tukey’s Honest Significant test; Suppl Fig S5d), namely Clostridiales, *Erysipelotrichaceae* and butyrate-producing bacteria from the *Lachnospiraceae* family. For instance, taxa such as *Barnesiella, Blautia, Butyrivibrio, Coprococcus, Dorea, Desulfovibrio, Hespellia, Lachnospira, Myroides* and *L-Ruminococcus* (all pertaining to the *Lachnospiraceae* family) were commonly found in less fit athletes in both discovery and validation sets. Contrastingly, individuals with improved cardiovascular fitness harbored a multitude of minor players, as observed in the discovery set. Although larger cohorts are required to clearly validate the relationship between athletic performance and gut microbiome, these data do, however, confirm the association between Firmicutes (notably *Lachnospiraceae* taxa) and cardiovascular fitness.

### Holo-omics: rare microbiomes with lower abundances of *Lachnospiraceae* taxa associated with improved cardiovascular fitness and points toward enhanced mitochondrial capacity

To characterize the microbiome-host crosstalk and identify molecular differences between the two types of cardiovascular outcomes in elite horses, we integrated multi-omic datasets from host and associated gut microorganisms through a multivariate matrix factorization approach using DIABLO (Data Integration Analysis for Biomarker discovery using Latent Components). To achieve this integrated perspective coined as holo-omics^34^, we combined pair host-centered omic and phenomics data with the shotgun metagenomics, fecal SCFAs composition and the concentrations of bacteria, anaerobic fungi and protozoa.

First, we observed strong covariation between the dominant phylotypes and the genetic functionalities derived from both KOs (*r*^2^ = 0.99) and CAZymes (*r*^2^ = 0.98). This clear correlation supports the added value of microbiome functionalities for status prediction rather than composition alone, as noted previously in human athletes ^14,35^. Concomitantly, the microbiome composition highly covaried with the mitochondrial transcriptome (*r*^2^ > 0.8) and the loads of fecal bacteria, anaerobic fungi and protozoa (*r*^2^ > 0.8; Fig 3a). Second, to add biological meaning to the predicted model, we investigated the relationship between the DIABLO-selected features with highest covariation (Suppl Fig S6a-c). The first latent variable of the predicted model indicated that athletes with higher cardiovascular fitness harbored a wide range of multi-kingdom and relatively low abundant species (Fig 3b). It included the facultative bacterial predator *Lysobacter*^36^, the health-promoting *Akkermansia*, which resides in the mucus layer of the gut and has been already reported in elite athletes ^39– 43^, along with anaerobic fungi (*Ophiocordyceps, Cryptococcus, Pseudogymnoascus, Trichoderma, Talaromyces)*, methanogens (*Methanothermobacter, Methanothrix*) and algae (*Emiliania* and *Porphyra*). Coupled with these rare yet highly active microbes, the first latent variable spanned CAZymes involved in the extraction of energy from recalcitrant polysaccharides and endogenous host glycans (GH99, GT10; Fig 3c). Conversely, less fit horses harbored higher amounts of core species, that is, dominant Firmicutes taxa (Clostridiales, *Erysipelotrichaceae* and multiple members of the family *Lachnospiraceae*), *Treponema* and *Prevotella*. Paired to it, the latter were characterized by functionally redundant enzymes with respect to lignocellulosic carbohydrate catabolizing machinery (GH8, GT36, GH51, GH28, GT2, GH5, GH3; Fig 3c). Although intestinal microbiota members belonging to the *Lachnospiraceae* family are known to produce significant amounts of acetate and butyrate ^37^, none of these SCFA were significantly increased in the feces or plasma of these athletes (*p* > 0.05) and the fecal pH remained unchanged (Suppl Table S15).

**Figure 3.**
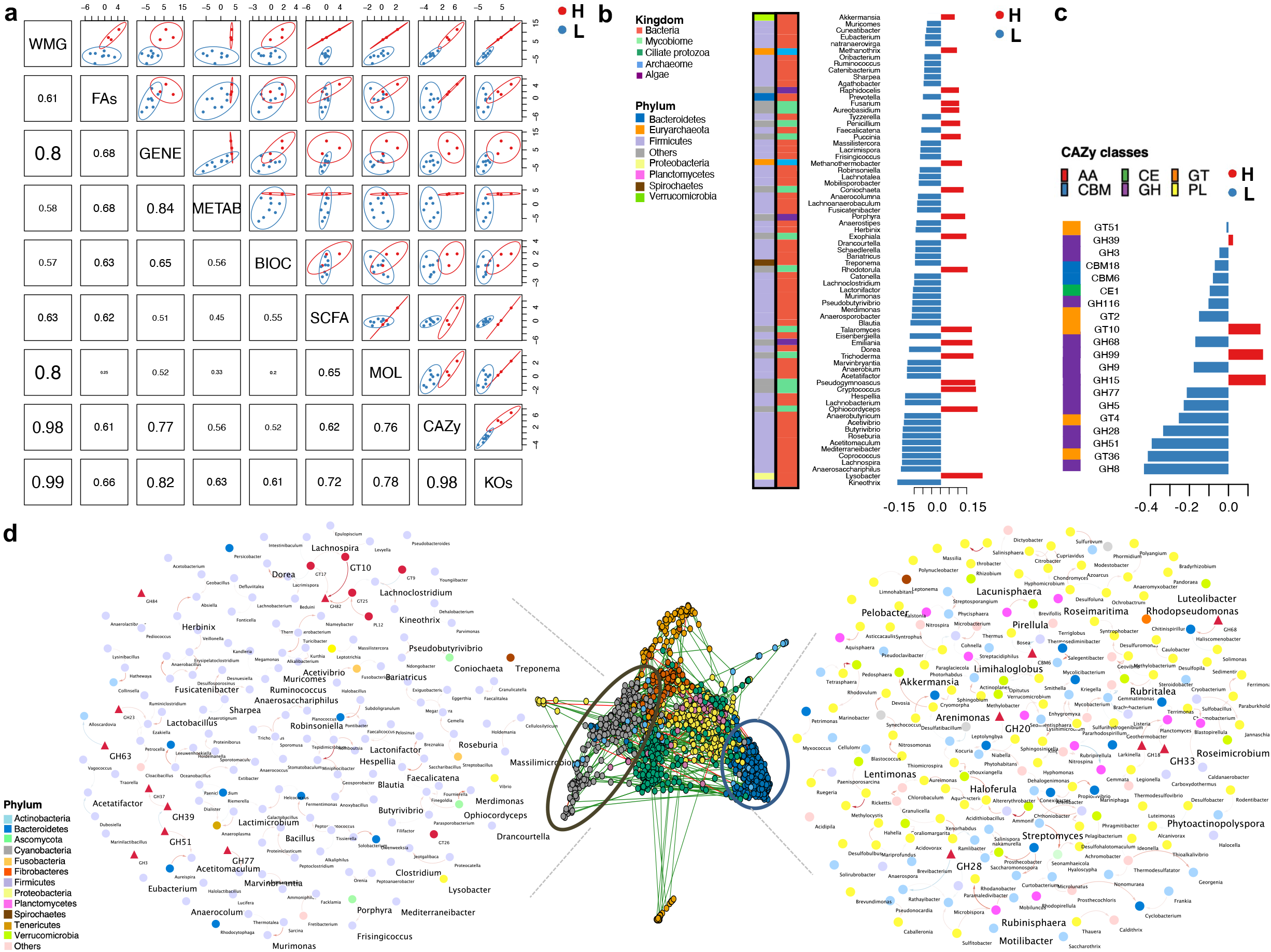
Sportomics: data integration supports the link between cardiovascular fitness and microbiome composition and functionality. (a) Matrix scatterplot showing the correlation between the first components related to each dataset in DIABLO according to the input design; (b) Microbial genera contributing to the separation along with component 1 of the microbiome dataset. Microbiome data are centered log-ratio-transformed and bar length indicates loading coefficient weight of selected phylotypes, ranked by importance, bottom to top. Columns on the left depict the kingdom and phylum of each discriminant phylotype; (c) CAZymes contributing to separation along with component 1 of (d). CAZymes profiles are log-transformed median-scaled values. Bar length indicates loading coefficient weight of selected CAZymes, ranked by importance, bottom to top. In all cases, colors indicate community classification, the community type 1 (red color) and community type 2 (blue color). Column in the left depict the CAZy class; (d) Co-occurrence network analysis of dominant phylotypes and carbohydrate-active enzymes (CAZy) classes datasets using sparse inverse covariance estimation for ecological association inference (SPIEC-EASI). Louvain clustering was able to generate 12 feature co-occurrence modules. The two extreme assortative modules are depicted in detail using Cytoscape. A positive correlation between nodes is indicated by red connecting lines, negative correlation by blue. Species and CAZymes features are denoted by a circle or triangle, respectively. Nodes are colored by phyla. Features with higher text size are those revealed as discriminant along with component 1 by the MixOmics approach. Edge width corresponds to the strength of the association between features.

An impairment rather than improvement of metabolic flexibility in the less fit individuals was supported by the reduced expression of genes in β-oxidation (*ECI1, SCP2, ACLY*), the electron transport chain (*TMEM242, NDFB4, TMEM126B, NDUFV3, NDUFA1, NDUFA10, SURF1, NDUFV1, DLD)*, Ca^2+^ translocation *(PMPCA, VDAC2, PHB)*, protein *(MRPL2, PPA1, MRPL49, MRPL17, VDAC2, TARS2, MRPL24, FBXL4, MRPS18C, PSTK)*, mitophagy (*TOMM40*) and mitochondrial biogenesis (*SSBP1, ACSS2*) (Suppl Fig S6d). The fact that adipose tissue lipolysis likely exceeded uptake and oxidation mitochondrial capacity in less fit individuals was confirmed by reduced concentrations of glucose (*p* = 0.0484, Wilcoxon rank-sum test), increased accumulation of long-chain acyl-carnitines (*i*.*e*., oleoyl carnitine, *p* = 0.0484; hydroxy oleoyl carnitine, *p* = 0.0242, Wilcoxon rank-sum test) and a tendency for augmented non-esterified fatty acids in plasma (*p* = 0.0848, Wilcoxon rank-sum test, see Suppl information). Additionally, less fit individuals showed depletion of metagenomic KOs involved in the mitochondrial biogenesis (K03593, K07152) and energy resilience (peroxisome proliferator-activated receptor (PPAR); K00029, K01596, K01897; see Suppl information). This could consequently decrease fatty acids oxidation but also increase glucose catabolism and progressively impede longer running times.

It is worth noting that the ciliate protozoal biomass, at up to about 18% of the biomass (∼10^9^ cells/g of stool), was representative of more fit individuals (Suppl Fig S6e). The main observed genera were *Stentor, Stylonychia, Pramecium* and *Tetrahymensa*, although their abundance and composition were much more variable than bacteria and their role is not well understood.

Altogether, less diverse microbial ecosystems dominated by few Firmicutes-derived phylum (mainly *Lachnospiraceae*) appear to set an upper-bound to the host metabolic response to exercise because of reduced mitochondrial energy production and biogenesis that ultimately constrain aerobic ATP production and extended cardiovascular fitness.

### Frenemies: *Lachnospiraceae* and rare phylotypes in athletes

To gain insight into the co-occurrence and co-exclusion relationships between multi-kingdom microbial genera, including organizational features that may contribute to host adaptation to exertion, we applied an inverse covariance estimation for ecological association inference between microbiome (at the genus level) and its metabolic potential, regardless of the cardiovascular fitness. This approach identified 12 modules. Among them, we uncovered two extreme assortative modules which were characterized by strong within microbe-microbe or microbe-functions interactions (Fig 3d) and recalled the features identified using DIABLO. The first module was mostly characterized by bacterial interactions within commensals from the Firmicutes phylum (mostly from *Lachnospiraceae* family) sharing similar phylogenetic and functional properties, along with CAZy families that can target the substrate of plant structural polysaccharides (GH3, GH39, GH51, GH82, GH84). On the other hand, the second extreme network encompassed widespread yet minor bacteria from the Proteobacteria, Actinobacteria, Planctomycetes, Verrucomicrobia (including *Akkermansia* sp.) and rare phyla, together with CAZymes active on degradation of complex structure of plant cell-wall materials (GH28) and host glycans (GH20, GH18, GH33; Fig 3d).

These results suggested again that ecosystems enriched in rare microorganisms are functionally different from Firmicutes dominant ones and cluster into segregated and distinct communities reflecting an ecological or evolutionary selective advantage to the microbiome.

## DISCUSSION

The current study presents the first horse gut microbiome gene catalog and its association with endurance performance. We have generated a catalog representing over 25 million non-redundant genes, expanding the current state of diversity for the equine gut microbiome. The building of this gene catalog has also widened twenty-fold the number of genera known to reside in the gastrointestinal tract of horses ^12,38,39^, uncovering an unprecedented number of prokaryota and eukaryota species mainly coming from the Ascomycota, Ciliophora, Basidiomycota, Chytridiomycota, Evosea and Apicomplexa phyla. Interestingly, this catalog captured a wide array of specific functions, suggesting that athlete gut microbiomes possess functional capacities primed for a greater ability to exploit energy from dietary, microbial and host resources and tissue repair as previously posited ^22^. Moreover, we identified 372 MAGs, most of which appear to be novel species. Although much of the gut microbiome likely represented unobserved diversity as our samples contained a significant proportion of unclassified sequences, the availability of so many novel genes and MAGs represents a significant step forward in understanding the composition and function of the horse gut microbiome.

Along with fatigue resistance, cardiovascular fitness is a key indicator of endurance performance in human athletes ^5^. Comparably, in horses, the cardiovascular capacity based on heart rate, heart recovery time after exercise and average speed across the race is considered to be a good indicator of the degree of peripheral and central fatigue during endurance exercise and thus, athletic performance ^40^. In our cohort of elite horses, ∼40% of the variation in the gut microbiome composition was accounted for by this parameter in absence of significant variation in dietary intake and across homogeneous genetic backgrounds. This finding was validated in an independent cohort of elite horses and echoes the association found between the cardiovascular fitness of marathon runners and their microbiota (∼ 22% of explained variance ^15^). Taking a closer look, less fit individuals were associated with Firmicutes taxa and particularly, the dominance of tightly related *Lachnospiraceae* spp. which are able to break down plant polysaccharides easily available in the gut. While this family is abundant in the adult human ^41^ and horse gut microbiome ^11,42,43^; its abundance can be rapidly altered by changes in diet ^44^. Therefore, in light of these findings, nutritional interventions to reduce *Lachnospiraceae* taxa abundance and functions while creating more space for rare species will be likely required to increase microbiome diversity and athletic performance. Some possible nutritional interventions could include probiotics and dietary fibers with higher specificity (*i*.*e*., accessible and fermentable by a limited range of microbes). On the other hand, a recent study proposes that fecal microbiome transplantation ^6^ as a means to increase exercise performance in athletes, raising the possibility of fecal modulation as a way to gain an athletic advantage.

Interestingly, the deep phenomics applied to these horses highlighted the microbiome-mitochondria axis as one of the most effective ways to modulate the cardiovascular capacity. Metagenomic and mitochondrial genes involved in the mitochondrial biogenesis and energy resilience were all simultaneously upregulated in more fit individuals, suggesting improved exercise economy and fuel sparing during endurance. Additionally, their gut microbiomes, characterized by a greater *α*-diversity and a vast range of rare genera, showed highly functional capabilities, spanning many aspects of breaking down plant polysaccharides and animal glycans such as glycosaminoglycan substrates (*i*.*e*., mucins, hyaluronan, heparin and chondroitin). Conceptually, individuals with improved cardiovascular fitness shifted the burden of microbiome nutritional support to host mucus glycans, hyaluronic acid and other glycoproteins from the intestinal environment and thus offer complementary or unique metabolic pathways to enhance the mitochondrial functioning and meet the high energy needs during exertion. Thus, it is speculative but not implausible that rare species (other than *Akkermansia*) might be able to degrade and consume mucus-like glycoproteins that reside near the gut mucosa, which allows them to influence the host adaptation to endurance exercise. In endurance horses, the consumption and catabolism of N-acetyl moieties of glycoproteins during intense exercise has already been observed ^45^. Yet, the role of the bacterial predator *Lysobacter*, the one with highest discriminative power in our DIABLO model, is not well understood; however, ecosystems enriched with predatory bacteria are known to be metabolically more active than other ecosystems and that they have important roles in regulating nutrient fluxes in microbial food webs ^52^.

The PPAR pathway might be the main mechanism through which SCFAs and secondary microbial metabolites from glycans and protein degradation in the lumen engage in multiple converging pathways to regulate mitochondrial functions in different tissues, including the heart. For example, PPAR-α is highly expressed in heart tissue where high levels of mitochondrial fatty acid oxidation occur ^53^. In line with the increased PPAR metagenomic pathway, more fit athletes showed increased expression of mitochondrial-related genes belonging to energy mitochondrial metabolism and biogenesis, Ca^2+^ cytosolic transport as well as inflammation, all of which are necessary to improve aerobic work capacity, spare glycogen usage and reduce peripheral fatigue ^54^. Mitochondria are essential for the physiological activity of the cardiovascular system due to their crucial role in bioenergetic and anabolic metabolism and their regulation of intracellular Ca^2+^ fluxes, which contribute to cardiac muscle contraction ^55^. Even the slightest decrease in their efficiency can have a profound impact on cardiovascular capacity ^56^. Despite this data, the role the microbiome directly has in mitochondrial function and density during exercise has yet to be elucidated.

This study has revealed for the first time the enormous levels of untapped microbial diversity, biotic interactions and functional gene potential in the gut of horse athletes. Using this unprecedented catalog of genes, our findings suggest that the variability of the gut microbiome composition and functions were associated with cardiovascular fitness in two ways. First, less diverse microbial ecosystems comprising high amounts of *Lachnospiraceae* taxa showed lower expression of metagenomic and mitochondrial related-genes assicuated with mitochondrial energetic activity, thereby leading to reduced amounts of aerobic ATP and impaired cardiovascular function and thus reduced athletic performance. Second, ecosystems harboring a large range of rare phylotypes, including the promising probiotic *Akkermansia*, were found to be metabolically more active and thus offer complementary or unique metabolic pathways to enhance the physiological activity of the cardiovascular system via crosstalk between mitochondria in peripheral tissues. Functional studies of gut microbiome species that are intimately connected with mitochondria function will be instrumental for the development of novel dietary strategies toward optimized cardiovascular capacity and therefore athletic performance.

## METHODS

### Ethics approval

The study protocol was reviewed and approved by the local animal care and use committee (ComEth EnvA-Upec-ANSES, reference: 11-0041, dated July 12^th^ 2011) for horse study. All the protocols were conducted in accordance with EEC regulation (n° 2010/63/UE) governing the care and use of laboratory animals, which has been effective in France since the 1^st^ of January 2013. In all cases, the owners and riders provided their informed consent prior to the start of sampling procedures with the animals.

### Animals

Eleven pure-breed or half-breed Arabian horses (3 females, 1 male and 7geldings; mean ± SD age: 10 ± 1.69) trained for endurance were selected from a cohort previously used in our team ^17,57–59^. All equine athletes started their training for endurance competitions at the age of 4 and presented a similar training history, level of physical fitness and training environment. The 11 horses were selected due to the following these criteria: 1) enrollment in the same 160 km endurance category; 2) blood sample collection before and after the race; 3) feces collection before the race; 4) absence of gastrointestinal disorders during the four months prior to enrollment; 5) absence of antibiotic treatment during the four months prior to enrollment and absence of anthelmintic medication within 60 days before the race; and 6) a complete questionnaire about diet composition and intake.

Subject metadata, including morphometric characteristics and daily macronutrients diet intake records are depicted in Suppl Table S1. Daily nutrient intakes calculations are described elsewhere ^57^.

### Performance measurement

The endurance race was split into successive phases of ∼30 – 40 km. At the end of each phase, horses were checked by veterinarians (referred to as a vet gate). The heart recovery time was the primary criterion evaluated at the vet gate as it is shown to be a remarkable complement to a physical assessment of an individual. At each vet gate, the heart rate was measured by the riders and a veterinarian using a heart rate meter and a stethoscope, respectively. Any horse deemed unfit to continue (due to a heart rate above 64 bpm after 20 min of recovery) was immediately withdrawn from the event. It should be noted that the time interval between arrival at the vet gate and the time needed to decrease the heart rate below 64 bpm was counted as part of the overall riding time. Therefore, the cardiac recovery time was calculated as the difference between the arrival time (at the end of the phase) and the time of veterinary inspection (referred to as the “time in” by the FEI endurance rules). The average speed of each successive phase was calculated at the vet gate.

Changes in these three variables during endurance events have shown to predict whether a horse is aerobically fit or not ^40^. In an attempt to estimate cardiovascular capacity, which is linked to performance capability and achievement, we consider all these variables together. Therefore, these three variables were first scaled through a *Z-*score, that is, the number of standard deviation units a horse’s score is below or above the average score. Such a computation creates a unitless score that is no longer related to the original units of analysis (*i*.*e*., minutes, beats, Km/h) as it measures the number of standard deviation units and therefore can more readily be used for comparisons. A composite based on such Z-scores was then created to estimate cardiovascular fitness. Specifically, the *“composite”* function (multicon R package, v.1.6) was used to create a unit-weighted composite of the three variables listed above.

### Transcriptomic microarray data production, pre-preprocessing and analysis

The transcriptome microarray data production, pre-processing and analysis is depicted in Plancade et al. (20219) and ^17^. Briefly, blood samples for RNA extraction were collected from each animal at T0 and T1 using Tempus Blood RNA tubes (Thermo Fisher).

Total RNAs were then isolated using the Preserved Blood RNA Purification Kit I (Norgen Biotek Corp., Ontario, Canada), according to the manufacturer’s instructions. Transcriptome profiling was performed using an Agilent 4×44K horse custom microarray (Agilent Technologies, AMADID 044466). All of the steps were conducted as described previously ^60,61^. We refer to our previous work for more details on the pre-processing, normalization and the application of linear models ^17^. Given our interest in understanding the role played by mitochondria during exercise, the set of 801 differentially expressed mitochondrial genes reported by our team ^17^ was selected for the downstream steps of analysis (Suppl Table S8).

### Proton magnetic resonance (^1^H NMR) metabolite analysis in plasma

As described elsewhere ^57,58^, the plasma metabolic phenotype of endurance horses was obtained from ^1^H NMR spectra at 600 MHz. Blood was collected from each horse the day before the event and within 30 minutes from the end of the endurance race using sodium fluoride and oxalate tubes in order to inhibit further glycolysis that may increase lactate levels after sampling. The ^1^H NMR spectra were acquired at 500 MHz with an AVANCE III (Bruker, Wissembourg, France) equipped with a 5 mm reversed QXI Z-gradient high-resolution probe. Further details on sample preparation, data acquisition, data quality control, spectroscopic data pre-processing and data pre-processing including bin alignment, normalization, scaling and centering are broadly discussed elsewhere ^62^. Details on metabolite identification are described in our previous work ^17,57^.

### Biochemical assay data production

Blood samples for biochemical assays were collected before and after the race using 10 mL BD Vacutainer EDTA tubes (Becton Dickinson, Franklin Lakes, NJ, USA). As detailed in ^57^, after clotting the tubes were centrifuged and the harvested serum was stored at 4 °C until analysis. Sera were assayed for total bilirubin, conjugated bilirubin, total protein, creatinine, creatine kinase, β-hydroxybutyrate, aspartate transaminase (ASAT), γ-glutamyltransferase and serum amyloid A levels on a RX Imola analyzer (Randox, Crumlin, UK).

### Blood acylcarnitine profiling

The serum acylcarnitine profiles, as a proxy for mitochondrial β-oxidation, were produced and analyzed as described elsewhere ^59^. Briefly, blood samples were collected in plain tubes prior to and within 30 minutes of the end of the ride. After clotting, the tubes were centrifuged and the harvested serum was stored at 4 °C for no more than 48 hours and subsequently stored at -80 °C. Free carnitine and a total of 27 acylcarnitines in plasma were analyzed as their butyl ester derivatives by electrospray tandem mass spectrometry (ESI-MS-MS) in the positive mode and detected on a triple quadrupole mass spectrometer (Xevo TQ-S Waters, Milford, MA, USA) using deuterated water.

### Fecal measurements: SCFA, DNA extraction and microorganism concentrations

Fresh fecal samples were obtained while monitoring the horses before the race. One fecal sample from each animal was collected off the ground immediately after defecation as described previously ^57,63^ and three aliquots (200 mg) were prepared. Since most of the horses experienced dehydration after the race, the gastrointestinal emptying was significantly delayed and therefore it was not possible to recover the feces immediately after the race. Aliquots for SCFA analysis and DNA extraction were snap-frozen.

SCFAs levels were determined by gas chromatography using the method described elsewhere ^64^.

Total DNA extraction from the 11 samples was performed as previously described ^17^. Briefly, DNA was extracted from ∼200 mg of fecal material using the EZNA Stool DNA Kit (Omega Bio-Tek, Norcross, Georgia, USA) and following the manufacturer’s instructions. DNA was then quantified using a Qubit and a dsDNA HS assay kit (Thermo Fisher).

As detailed in our previous studies ^17,57^, concentrations of bacteria, anaerobic fungi and protozoa in fecal samples were quantified by qPCR using a QuantStudio 12K Flex platform (Thermo Fisher Scientific, Waltham, USA). Primers for real-time amplification of bacteria (FOR: 5’-CAGCMGCCGCGGTAANWC-3’; REV: 5’-CCGTCAATTCMTTTRAGTTT-3’), anaerobic fungi (FOR: 5’-TCCTACCCTTTGTGAATTTG-3’; REV: 5’-CTGCGTTCTTCATCGTTGCG-3’) and protozoa (FOR: 5’-GCTTTCGWTGGTAGTGTATT-3’; REV: 5’-CTTGCCCTCYAATCGTWCT-3’). Details of standard dilutions series, the thermal cycling conditions and the estimation of the number of copies are detailed in ^57^ and ^17^.

### Fecal microbiota: V3–V4 16S rRNA gene sequencing and data pre-processing

Detailed description of the DNA isolation process, V3-V4 16S rRNA gene sequencing-PCR amplification is presented by our group ^11,12,17,57,63,65,66^.

The Divisive Amplicon Denoising Algorithm (DADA) was implemented using the DADA2 plug-in for QIIME 2 (v. 2021.2) to perform quality filtering and chimera removal and to construct a feature table consisting of read abundance per amplicon sequence variant (ASV) by sample ^67^. Taxonomic assignments were given to ASVs by importing Greengenes 16S rRNA Database (release 13.8) to QIIME 2 and classifying representative ASVs using the naive Bayes classifier plug-in ^68^. The phyloseq (v.1.36.0) ^69^, vegan (v.2.5.7) ^70^ and microbiome (v.1.14.0) packages were used in R (v.4.1.0) for the downstream steps of analysis. A total of 364,026 high-quality sequence reads were recovered for the 11 horses of the study (mean per subject: 33,093 ±17,437, range: 12,052 – 62,670). Reads were clustered into 5,412 chimera- and singleton-filtered ASVs at 99% sequence similarity. The ASV taxonomic assignments and ASV counts for each individual are presented in the Suppl Table S4).

### Fecal metagenome: Shotgun sequencing data production and analysis

Metagenomic sequencing was performed using the same DNA extractions. For each individual, a paired-end metagenomic library was prepared from 100 ng of DNA using the DNA PCR free Library Prep Kit (Illumina, San Diego, CA, USA) and size selected at about 400 bp. The pooled indexed library was sequenced in an Illumina HiSeq3000 using a paired-end read length of 2×150 pb with the Illumina HiSeq3000 Reagent Kits at the PLaGe facility (INRAe, Toulouse).

### MAG assembly and annotation

Raw metagenomics reads were quality-trimmed, assembled, binned and annotated using the ATLAS pipeline, v. 2.4.4 ^71^. In short, using tools from the BBmap suite v.37.99 ^72^, reads were quality trimmed and contamination from the horse genome were filtered out (available at NCBI sequence archive with the accession number GCA_002863925.1; Equus_caballus.EquCab3.0). Reads were error corrected and merged before assembly with metaSPAdes v.3.13.1 ^73^. Since a high diversity between individuals was described through 16S rRNA amplicon analysis, we first assembled each sample independently. QUAST 5.0.2 ^74^ was used to evaluate the quality of each sample assembly. Contigs from single samples were binned using MetaBAT 2 (v.2.14) ^75^ and Maxbin 2.0 v.2.2.7 ^76^ and their predictions were combined using DAS Tool v.1.1.2-1 ^77^.

The quality of the metagenome-assembled genomes (MAGs) was then assessed using checkM v.1.1.3 ^78^. The predicted MAGs presented at least 50% completeness and < 10% contamination. Because the same MAG may be identified in multiple samples, dRep v.2.2.2 ^79^ was used to obtain a non-redundant set of MAGs by clustering genomes to a defined average nucleotide identity (ANI, default 0.95) and returning the representative with the highest dRep score in each cluster. dRep first filtered genomes based on genome size (default > 5,000 bp) and quality (default > 50% completeness, < 10% contamination). MAGs were scored on the basis of completeness, contamination, genome size and contig N50, with only the highest scoring MAG from each secondary cluster being retained as the winning genome in the dereplicated set. The abundance of each MAG was then quantified across samples by mapping the reads to the non-redundant MAGs and determining the median coverage in 1 Kb windows along each genome.

For the taxonomic annotation, ATLAS predicted the genes of each MAG sequence using Prodigal v.2.6.3 ^80^ with default parameters. Robust taxonomic annotation was assigned to bins according to the genome taxonomy database (GTDB-tk ^81^) release 95, v.5.0 (July 17, 2020). As such, GTDB-Tk taxonomy names are used throughout this paper. In addition, MAG phylogenetic trees were built based on markers from GTDB-Tk and CheckM and visualized using ggtree (v.3.0.2) in R package.

To assess the contribution of the constructed MAGs to the functional potential of the gut microbiome, the predicted gene and proteins extracted by Prodigal during the CheckM pipeline were compared to the EggNOG database 5.0 using eggnog-mapper (v2.0.1). From this output, KEGG annotation (Kyoto Encyclopedia of Genes and Genomes) and CAZymes annotation (Carbohydrate-active Enzyme) were extracted. Since the detection of KOs and CAZymes families are likely to be influenced by sequencing depth, we first normalized their abundance relative to the abundance of the MAG they derived from. Pathways attributed to each KO were annotated from the KEGG Database (downloaded 23-October-2021; https://www.genome.jp/brite/ko00001).

The uniqueness of our predicted MAG catalog was confirmed by dereplicating them with the 121 MAGs produced by (Gilroy et al., 2021) and 3 reported by (Youngblut et al., 2020) using dRep v.3.2.0 ^79^. dRep performed pairwise genomic comparisons by sequentially applying an estimation of genome distance and an accurate measure of average nucleotide identity. The visualization and comparison of highly similar genomes were performed using the CGView family of tools (http://wishart.biology.ualberta.ca/cgview/).

### Construction of the integrated gene catalog

The establishment and assessment of the quality and representation of the microbiome gene catalog was performed through the metagenomic ATLAS pipeline (v.2.4.4) ^71^. As described above, we first assembled the clean reads into longer contigs.

Genes were predicted by Prodigal v.2.6.3 and then clustered using linclust ^82^ to generate a non-redundant gene catalog. Redundant genes were removed (≥ 95% identity and ≥ 90% overlap) with linclust. The quantification of genes per sample was done through the “*combine_gene_coverages*” function in the ATLAS workflow, which aligned the high-quality clean reads to the gene catalog. Taxonomic and function annotations were done based on the EggNOG database 5.0 using eggnog-mapper (v.2.0.1). From these, the eggNOG numbers corresponding to CAZymes based on homology searches to the CAZyme database were retrieved. We used the derived eggNOG abundance matrix to obtain a CAZyme profile per sample. Similarly, KEGG annotation was retrieved from the EggNOG output. KEGG gene IDs were mapped to KEGG KOs and used to obtain the KEGG functional pathway hierarchy.

### Annotation of metagenome using Kaiju

The k-mer-based kaiju v. 1.8.0. (https://github.com/bioinformatics-centre/kaiju) ^20^ approach was used for microbial taxonomic profiling of the shotgun metagenomes. Paired reads after quality trimmed and decontamination from the horse genome were used and annotated against the NCBI *nr* reference database (released on May 25^th^ 2020) containing all proteins belonging to archaea, bacteria, eukaryota and virus for classification in Greedy run mode with -a greedy -e 3 allowing for maximum three mismatches. By default, Kaiju returned a “NA” if it could not find a taxonomic classification at certain ranks.

### Resistome

The high-quality clean paired reads were aligned to the ResFinder database (accessed March 2018, v.4.0) using bowtie2 (v.2.3.5). ResFinder is a manually curated database of horizontally acquired antimicrobial resistance (AMR) genes and contains many genes with numerous highly similar alleles (*i*.*e*., β-lactamases). To avoid random assignment of read pairs on these high-identity alleles, the database was clustered at 95% of identity level, over 200 bp using CDHIT-EST (options -G 0 -A 200 -d 0 -c 0.95 -T 6 -g 1) ^83^ and a reference sequence was attributed to each cluster. Two successive mappings were done: (i) a first mapping with standard parameters (bowtie2 --end-to-end --no-discordant --no-overlap --no-dovetail --no-unal) on the complete ResFinder database and (ii) a second mapping on the clustered database using the reads from the first mapping, with less stringent parameters (bowtie2 --local --score-min L,10,0.8). More than 99% of the reads from the first mapping correctly aligned on a cluster reference sequence in the second mapping.

Counts from the second mapping were normalized by computing the RPKM (reads per kilobase reference per million bacterial reads) value for each ResFinder reference sequence. The RPKM values were computed by dividing the mapping count on each reference with its gene length and the total number of bacterial read pairs for the samples and multiplying by 10^9^. A minimum of 20 mapped reads was considered to validate the presence of an AMR gene cluster.

### Biodiversity and richness analysis: α- and β-diversity

The microbiome R package allowed us to study global indicators of the gut ecosystem state, including measures of evenness, dominance, divergences and abundance. Comparison of the gut α-diversity indices between groups was performed by two-tailed Wilcoxon test (pairwise comparison). Benjamini-Hochberg multiple testing correction *p*□<□0.05 was set as the significance threshold for the comparisons between groups.

To estimate β-diversity, Bray-Curtis dissimilarity was calculated using the phyloseq R package. All samples were normalized using the “*rarefy_even_depth*” function in the phyloseq R package, which is implemented as an *ad hoc* means to normalize features that have resulted from libraries of widely differing sizes. The PerMANOVA test (a non-parametric method of multivariate analysis of variance based on pairwise distances) implemented in the “*adonis2”* function from the vegan R package allowed testing the global association between ecological or functional community structure and groups.

The core microbiome of individual samples was calculated using a detection threshold of 0.1% and a prevalence threshold of 95% in the microbiome R package.

### The inter-individual variations in the gut microbiome composition and function

The inter-individual variations in the gut microbiome composition and function were studied based on the conceptual framework of community types ^84^. According to this framework, the samples were clustered into bins based on their taxonomic similarity ^85^. Briefly, clustering was performed with PAM ^86^ using Bray-Curtis distance of the normalized feature counts. The optimal number of communities was chosen by the maximum average silhouette width, known as the silhouette coefficient (SC) ^87^.

### Inference and Analysis of SPIEC-EASI Microbiome Networks

The SParse InversE Covariance Estimation for Ecological Association Inference method (SPIEC-EASI) ^88^ was used to identify sub-populations (modules) of co-abundance and co-exclusion relationships between dominant phylotypes and CAZy classes abundances matrices. Specifically, the method allows microorganisms and functions to interact in a number of different ways, from bidirectional competition to mutualism or to not interact at all. The statistical method SPIEC-EASI comprises two steps, first a transformation for compositionality correction of the feature matrices and second an estimation of the interaction graph from the transformed data using sparse inverse covariance selection. The sparse graphical modeling framework was constructed using the “*spiec*.*easi*” function of the SpiecEasi package (v.1.1.1). The features were clustered using the method = mb, lambda.min.ratio = 1e^-5^, nlambda = 100, pulsar.params=list (thresh = 0.001). Regression coefficients from the SPIEC-EASI output were extracted and used as edge weights to generate a feature co-occurrence network R igraph package (v.1.2.6) and Cytoscape (v.3.8.2).

### Integrative statistical analysis

Data integration was carried out using several approaches and different combinations of data sets. Prior to the integration, we applied some additional pre-processing steps on our explanatory data sets. In particular, to eliminate intra-individual variability and focus on the respective differential signals between T1 and T0, we considered Δ values (T1–T0) for each of these data sets, namely biochemical assay data, metabolome data, acylcarnitine profiles and gene expression data, as previously described ^17^. For the transcriptome, we constructed a matrix of log-transformed expression values between T1 and T0 (*i*.*e*., the difference in log_2_- normalized expression between T1 and T0, equivalent to the log_2_ value of the T1/T0 ratio) for the differentially expressed mitochondrial-related genes (Suppl Table S8).

The integration of data was then performed using complementary methods and working with different data sets available, namely: (1) Δ values of mitochondrial-related genes; (2) Δ values of ^1^H NMR metabolites; (3) Δ values of the biochemical assay metabolites; (4) Δ values of plasmatic acylcarnitines; (5) the fecal SCFAs at T0; (6) the bacterial, ciliate protozoal and fungal loads at T0; (7) the dominant gut phylotypes at T0; (8) the CAZymes profiles at T0; (7) the KOs at T0 and the (8) athletic performance data.

As a first integration approach, a global non-metric multidimensional scaling (NMDS) ordination was used to extract and summarize the variation in microbiome composition using the “*metaMDS*” function in the vegan R package. To determine the number of dimensions for each NMDS, stress values were calculated.

The explanatory data sets were then fit to the ordination plots using the “*envfit*” function in the vegan R package ^89^ with 10,000 permutations. The effect size and significance of each covariate were determined and all of the *p*-values derived from the “*envfit*” function were Benjamini-Hochberg adjusted. Variation partitioning was performed using the “*varpart”* function in vegan in R. The “*varpart”* function uses linear constrained ordination to assess the shared and independent (partialling out the others) contributions (adjusted R^2^) of several covariates on microbiome composition variation.

As a second integrative approach, the N-integration algorithm DIABLO of the mixOmics R package (http://mixomics.org/, v6.12.2) was used. It is to be noted that, in the case of the N-integration algorithm DIABLO, the variables of all the data sets were also centered and scaled to unit variance prior to integration. In this case, the relationships existing among all data sets were studied by adding a further categorical variable, *i*.*e*., the cardiovascular fitness of horses. Horses that had poor cardiovascular fitness (*n* = 8) were compared to horses that had enhanced cardiovascular fitness (*n* = 3). DIABLO seeks to estimate latent components by modelling and maximizing the correlation between pairs of pre-specified datasets to unravel similar functional relationships between them ^90^. A full weighted design was considered. To predict the number of latent components and the number of discriminants, the “*block*.*splsda*” function was used. In both cases, the model was first fine-tuned using the leave-one-out cross-validation by splitting the data into training and testing. Then, classification error rates were calculated using balanced error rates (BERs) between the predicted latent variables with the centroid of the class labels using the “*max*.*dist*” function.

Additionally, the DESeq2 (v. 1.32.0) ^91^ R package was used to test for differential abundances analysis between groups for each independent omic dataset. DESeq2 assumes that counts can be modeled as a negative binomial distribution with a mean parameter, allowing for size factors and a dispersion parameter. Next to the group, the horse dependency was included in the generalized linear model. The *p*-values were adjusted for multiple testing using the Benjamini-Hochberg procedure. DESeq2 comparisons were run with the parameters fitType□=□“parametric” and sfType□=□“poscounts”.

### The validation cohort

The validation set consisted of 22 pure-breed or half-breed Arabian horses (12 females, 3 male and 7 geldings; age: 9.2 ± 1.27) not included in the experimental set to ensure that the observed effects were reproducible in a broader context (Suppl Table S13). Among the horses in the validation set, five animals were enrolled in a 160 km endurance competition, while 17 horses were enrolled in a 120 km race. The management practices throughout the endurance ride and the International Equestrian Federation (FEI) compulsory examinations, as well as the weather conditions, terrain difficulty and altitude were that of the experimental set. In fact, all the participants enrolled in the study (experimental and validation set) competed in the same event during October 2015 in Fontainebleau (France). The cardiovascular capacity was created as described in the “Performance measurement” section, that is, as a composite of post-exercise heart rate, cardiac recovery time and average speed during the race. Then after, the HIGH, MEDIUM and LOW groups were determined according to the interquartile range of the composite cardiovascular fitness values, where HIGH included individuals with cardiovascular fitness values above the 75^th^ percentile, LOW below the 25^th^ percentile and MEDIUM the individuals ranging in between.

## Supporting information

Metadata of horses recruited in the experiment

Sequencing data and assembly statistics for each sample and quality assessment for the gene catalog

Top 25% most dominant phylotypes sorted by their abundance and ubiquitousness found in more than half of the samples. Taxonomic profiling was done wit

Number of ASVs and relative abundance of genera detected by 16S rRNA gene sequencing

Abundance of the CAZymes, classes and their estimated mechanisms in the gene catalog

List of acquired antimicrobial resistance (AMR) genes observed in this study

Assembly statistics of the 372 metagenome assembled genomes (MAGs)

List of differentially expressed mitochondrial-related genes

Relative abundance of metabolites obtained from blood of the 11 horses under study collected before and after the endurance ride

Biochemical parameters obtained from the blood of the 11 horses under study collected before and after the endurance race

Summary of the significant correlations between environmental and host variables and microbial community beta diversity ordination on NMDS plots

Metadata of the horses recruited in the validation set

ASV taxonomic assignments and ASV counts for the 22 horses in the validation set based on 16S rRNA sequencing

Fecal pH and fecal short chain fatty acids measurements in the 11 horses as well as concentrations of bacteria, protozoa and fungi

- Comparison of differential abundance of microbial phylotypes between individuals with different cardiovascular capacity.

Comparison of differential abundance of KOs and their associated pathways between individuals with different cardiovascular fitness

Comparison of differential abundance of CAZymes between individuals with different cardiovascular capacity.

Distributions of CAZymes in the 372 MAGs

Distributions of KOs in the 372 MAGs

Comparison of differential abundance of MAGs between pathways between individuals with different cardiovascular capacity.

## Data Availability

The datasets presented in this study can be found in different online repositories. Microarray expression data are available in Gene Expression Omnibus (GEO) repository under the accession number GSE163767 (https://www.ncbi.nlm.nih.gov/geo/query/acc.cgi?acc=GSE163767). Metabolomic data are available in the NIH Common Fund’s Data Repository and Coordinating Center UrqK1489; (http://dev.metabolomicsworkbench.org:22222/data/DRCCMetadata.php?Mode=Study&StudyID=ST000945). The gut metagenome 16S rRNA targeted locus data are available in the DDBJ/EMBL/GenBank under the accession KBTQ00000000.1; (locus KBTQ01000000). The corresponding BioProject is PRJNA438436 and the accession numbers of the BioSamples included in here are SAMN08715729, SAMN08715728, SAMN08715727, SAMN08715725, SAMN08715723, SAMN08715721, SAMN08715719, SAMN08715718, SAMN08715714, SAMN08715713, SAMN08715710. The validation set data is available under the same BioProject ID. Moreover, the raw metagenomic sequence data of the 11 athletes reported in this paper have been deposited in the NCBI short read archive (SRA) under the same BioProject ID PRJNA438436. The temporary submission ID is SUB10812702. All metagenome assemblies and sequences of MAGs have been deposited in NCBI under the same BioProject ID PRJNA438436. The temporary submission ID is <u>SUB10812003</u>. All other data is available in the Supplementary Data and upon reasonable request to the corresponding author.

## ACKNOWLEDGEMENTS

We are grateful for Marine Beinat, Julie Rivière, Emmanuelle Rebours, Jordi Estellé, Caroline Morgenthaler and Arni Janssen for participating in the sample collection and organization during the project. We particularly thank Valentine Ballan and Fanny Blanc who helped isolate the blood RNA. We also thank Catherine Philippe for the SCFA analysis in feces, Diane Esquerré who prepared the libraries and performed the MiSeq sequencing at the GeT-PlaGe genomics core facility (INRAE, Toulouse, https://get.genotoul.fr/) and Estel Blasi, who created the picture of horses. Lastly, we are grateful for all the horse owners, riders and endurance race organizers who participated in the study.

This work was funded by grants from the *Fonds Eperon*, the *Institut Français du Cheval et de l’Equitation* (IFCE), the *Association du Cheval Arabe* (ACA) and the *Institute National de la Recherche pour l’Agriculture, l’alimentation et l’environnement* (INRAE).

## AUTHOR CONTRIBUTIONS

NM, CR and EB conceived the study. NM and GS wrote the manuscript. CM and OR optimized the assembly pipeline to create the gene catalog and MAGs repertoire. SL analyzed the resistome. NM performed the integrative and statistical analysis. SP prepared the metagenome sequencing libraries. LLM performed the metabolomic experiment and analyzed the metabolite peaks. CR and EB were in charge of managing the GenEndurance project and organizing the sample collection during the race. GS, EB, CR, CM, SL and LLM revised the manuscript and contributed to the interpretation of the data. All authors read and approved the final manuscript.

