## Supplementary figures and images for "The first horse gut microbiome gene catalog reveals that rare microbiome ensures better cardiovascular fitness in endurance horses"

### Suppl Fig S1

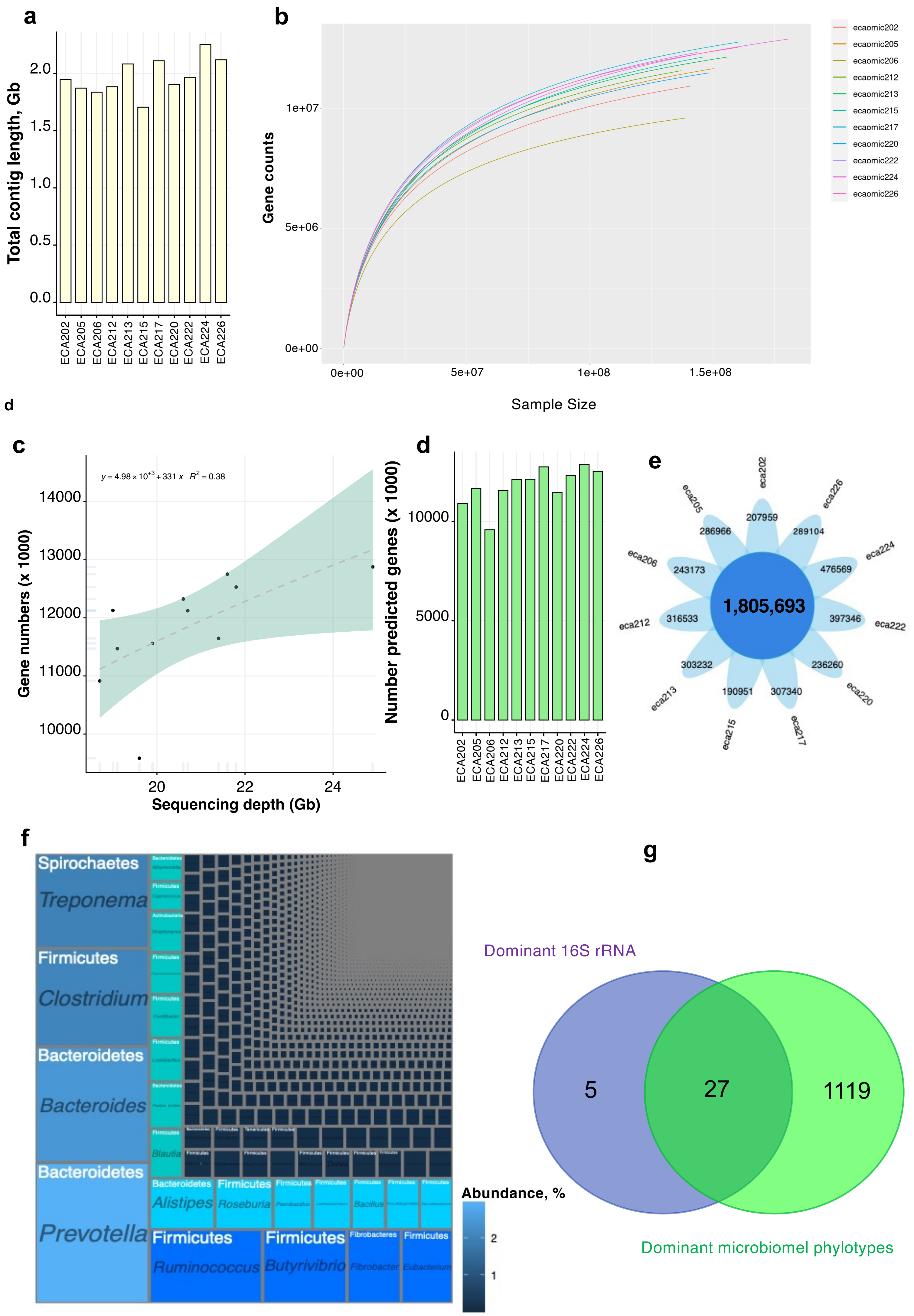

### Suppl Fig S2

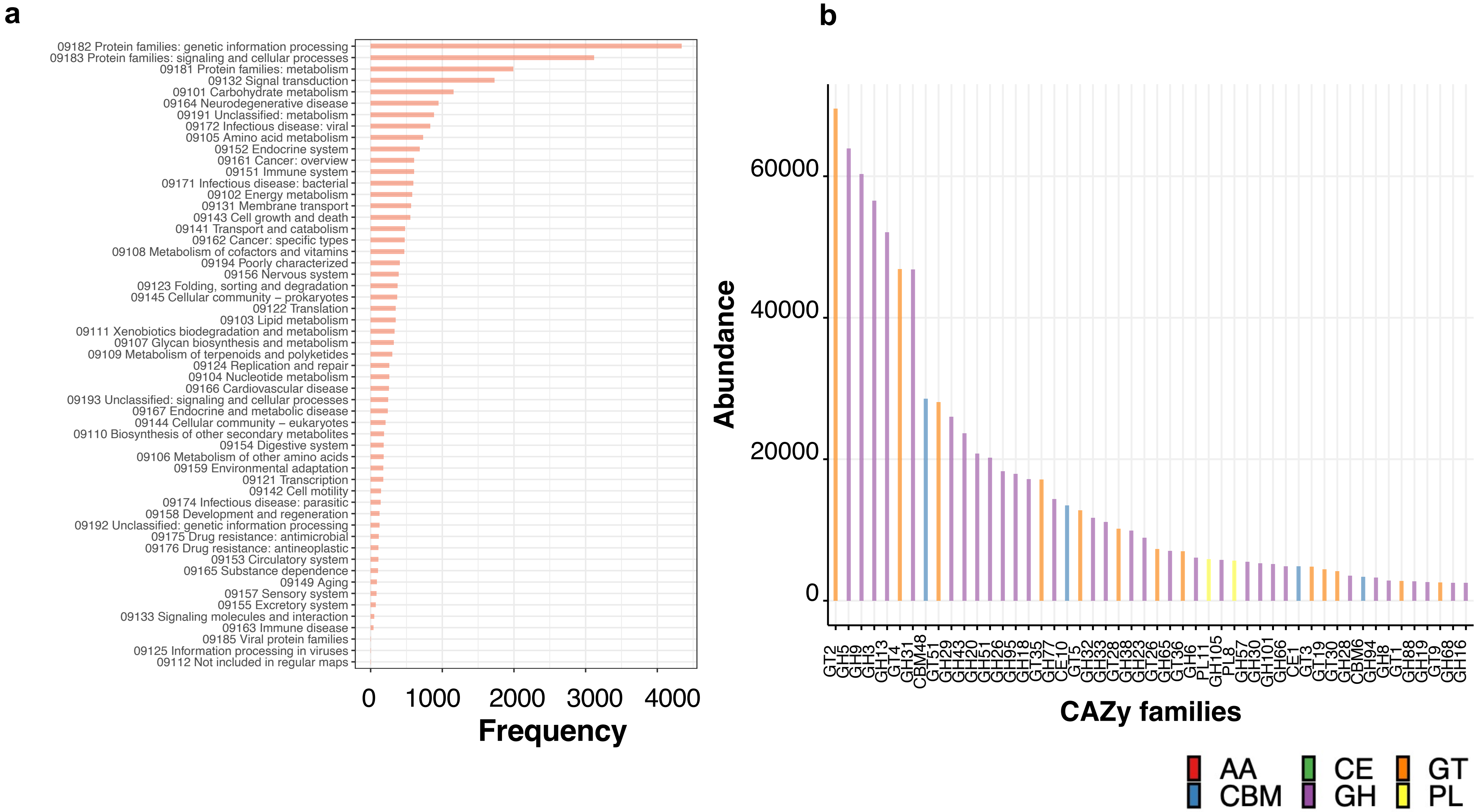

### Suppl Fig S3

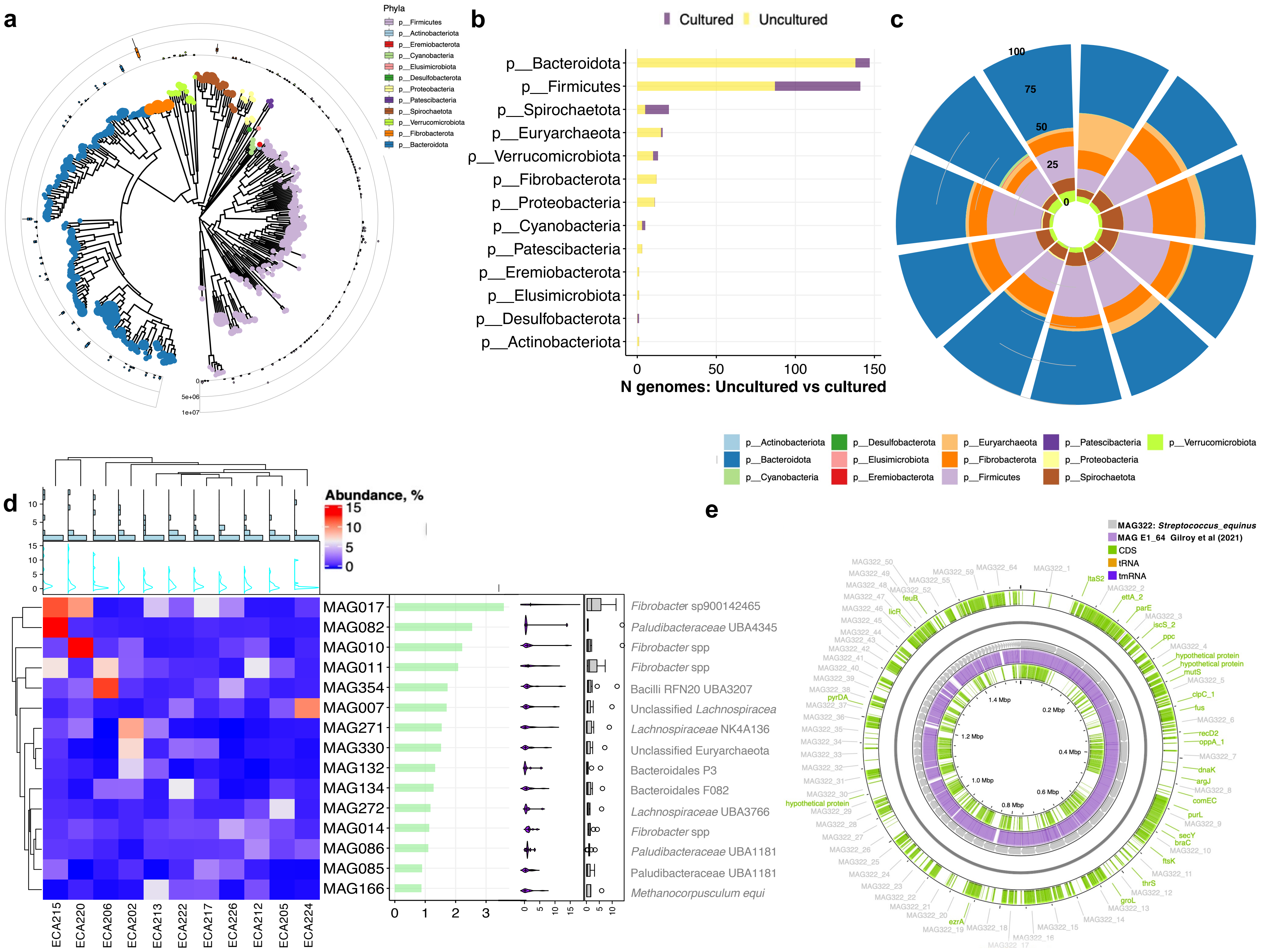

### Suppl Fig S4

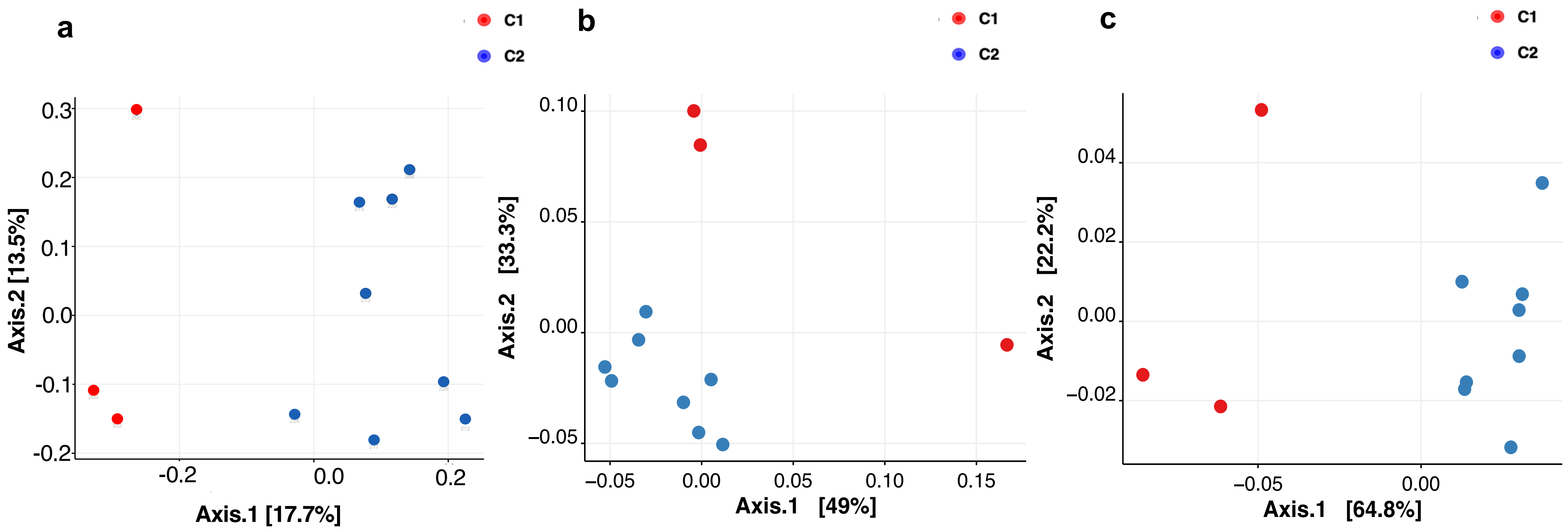

### Suppl Fig S5

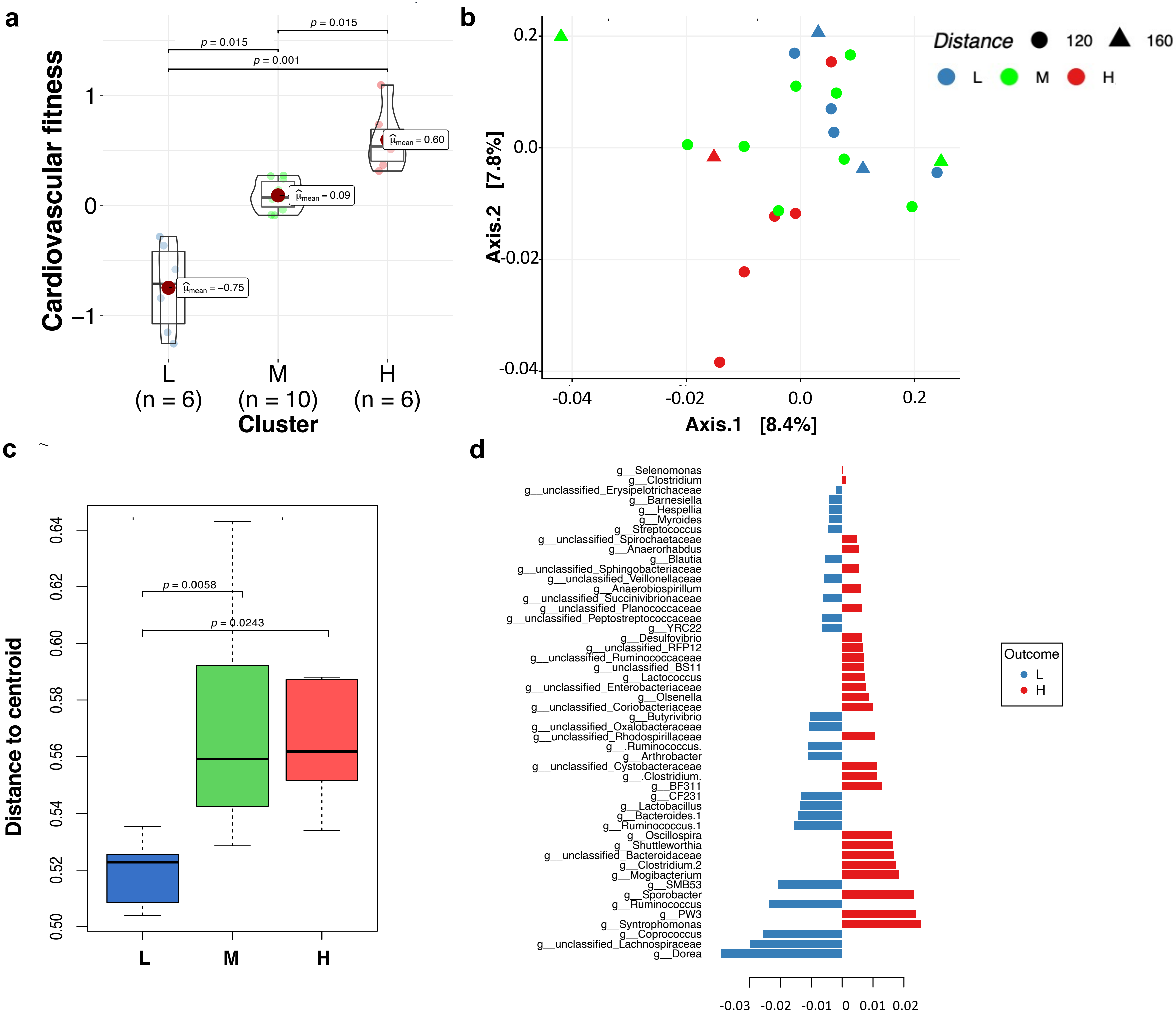

### Suppl Fig S6

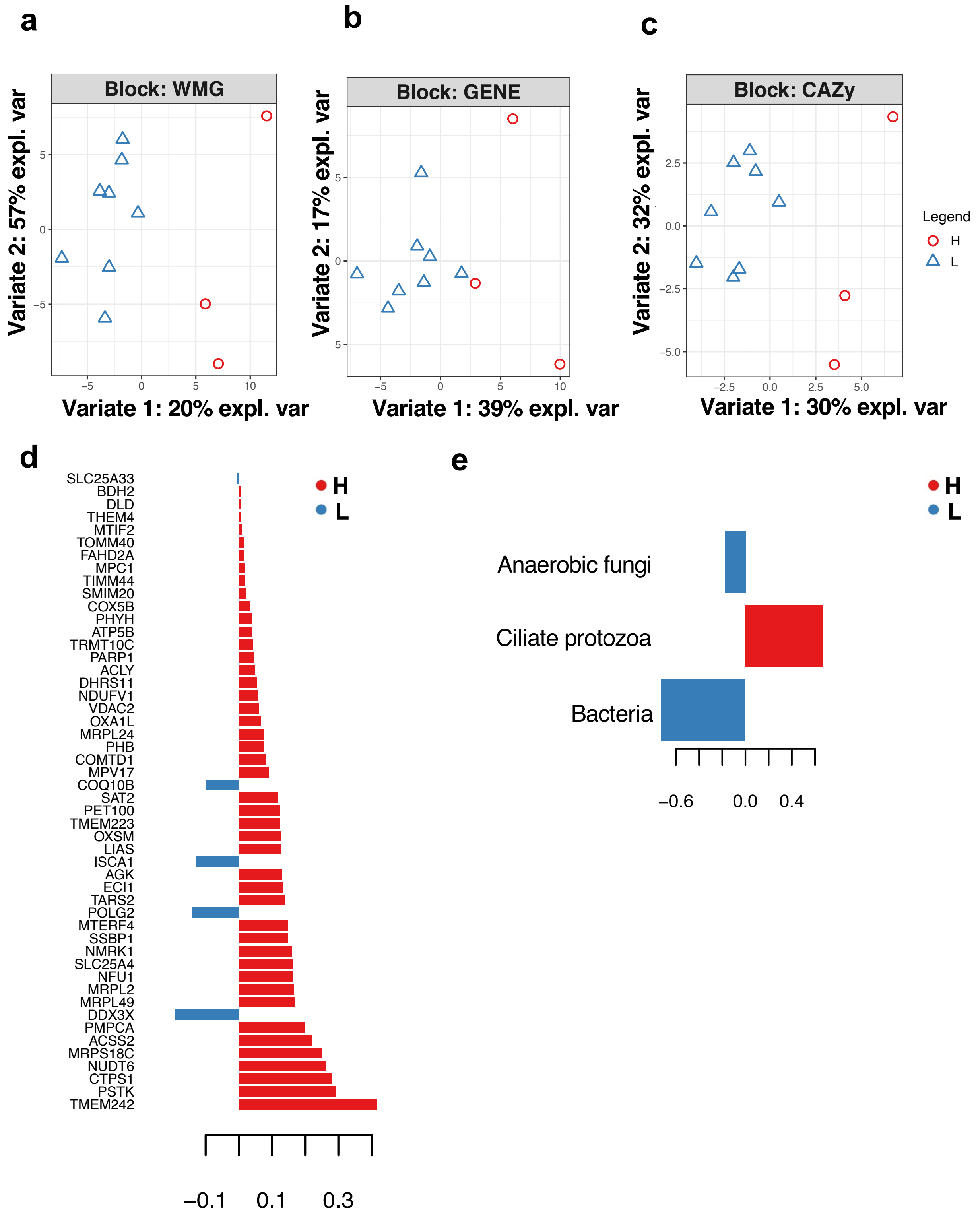

### Suppl Fig S7

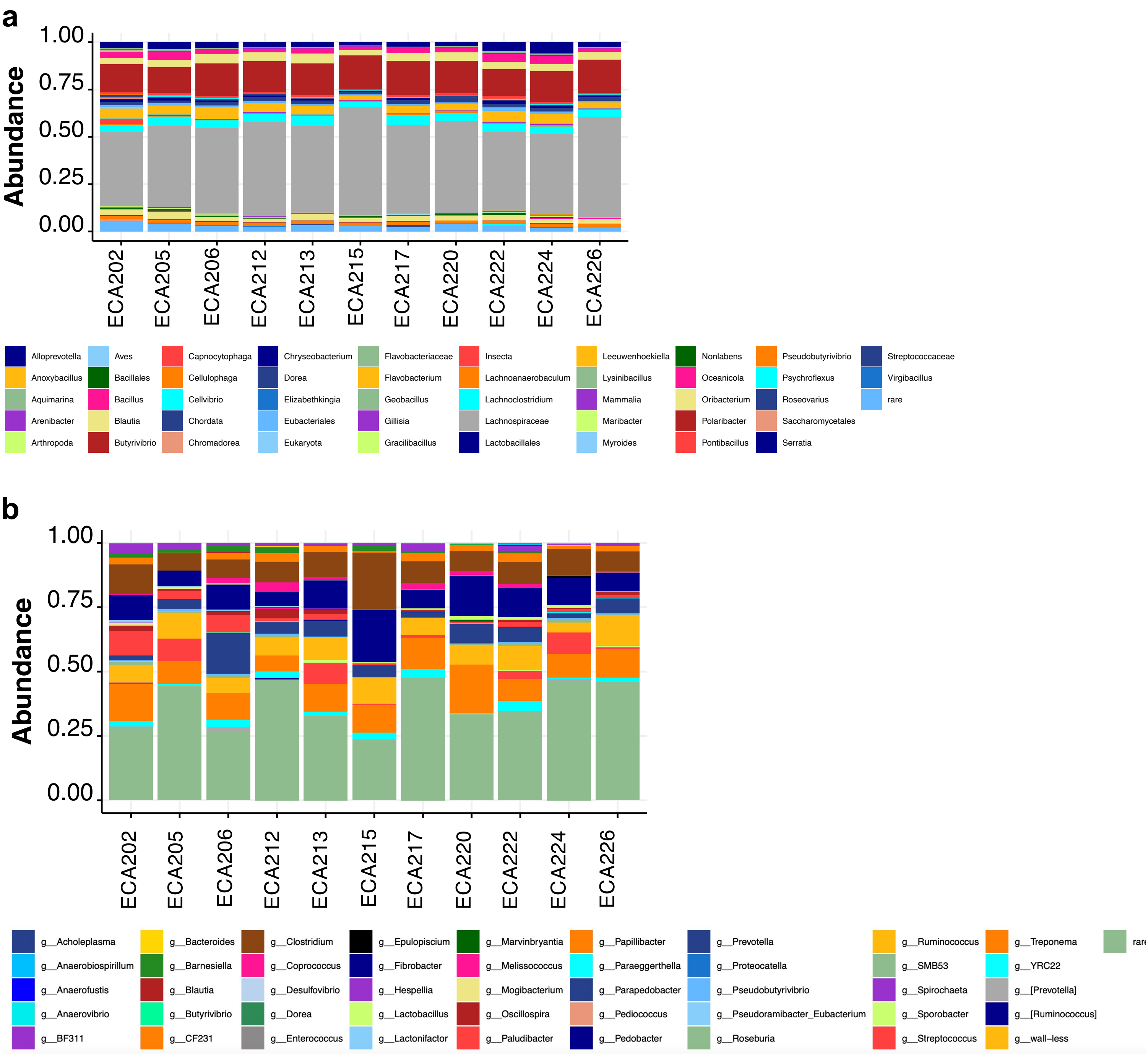

### Suppl Fig S8

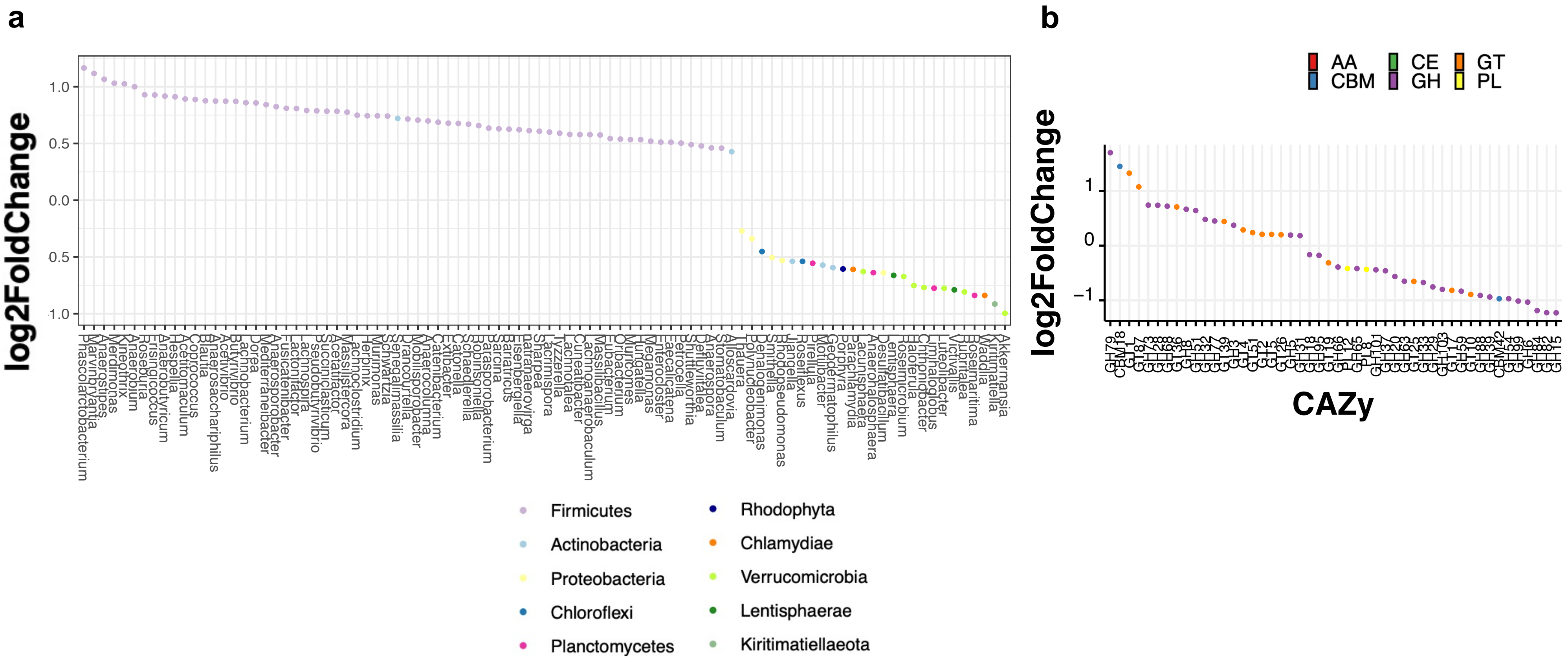

### Suppl Fig S9

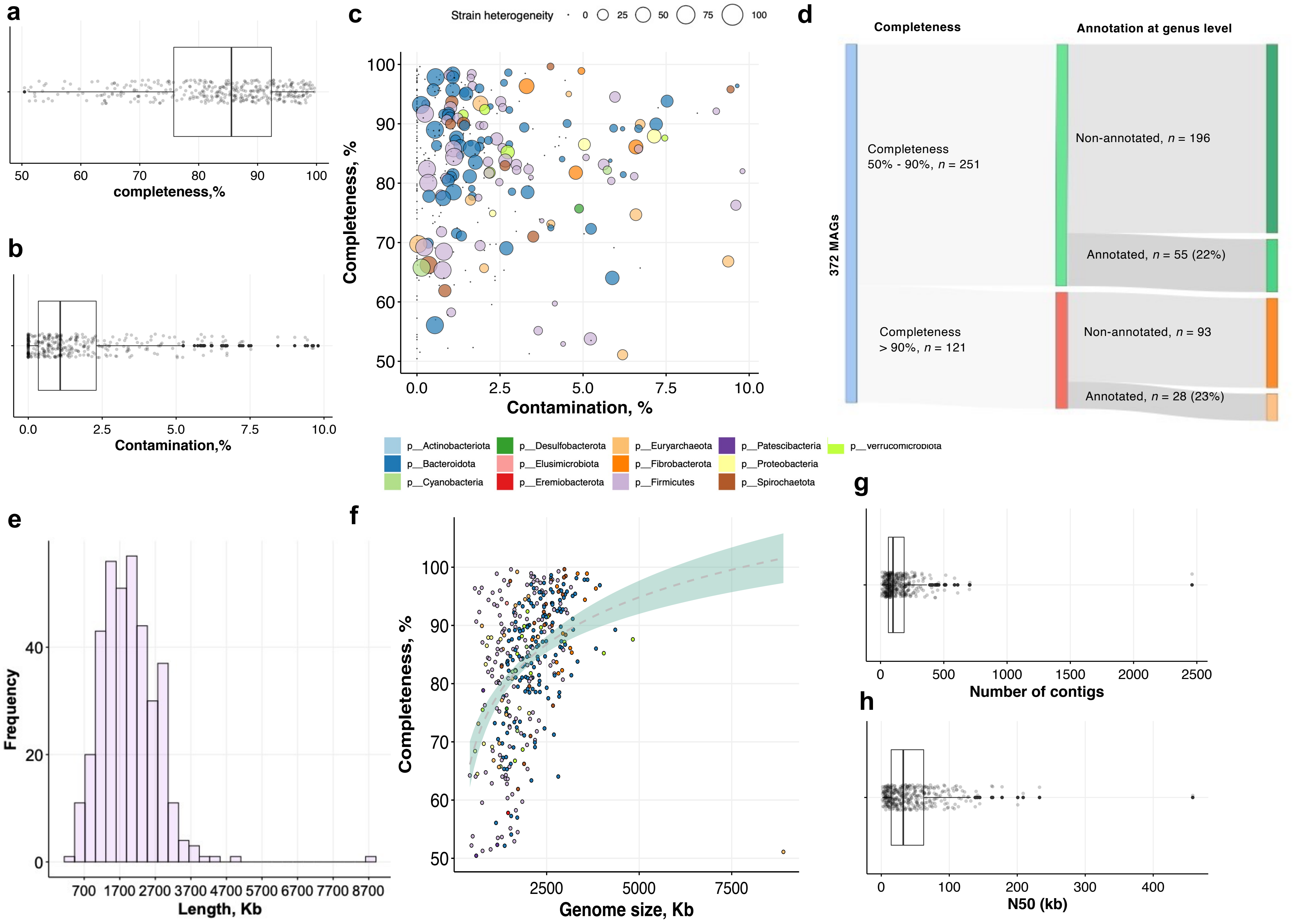

### Suppl Fig S10

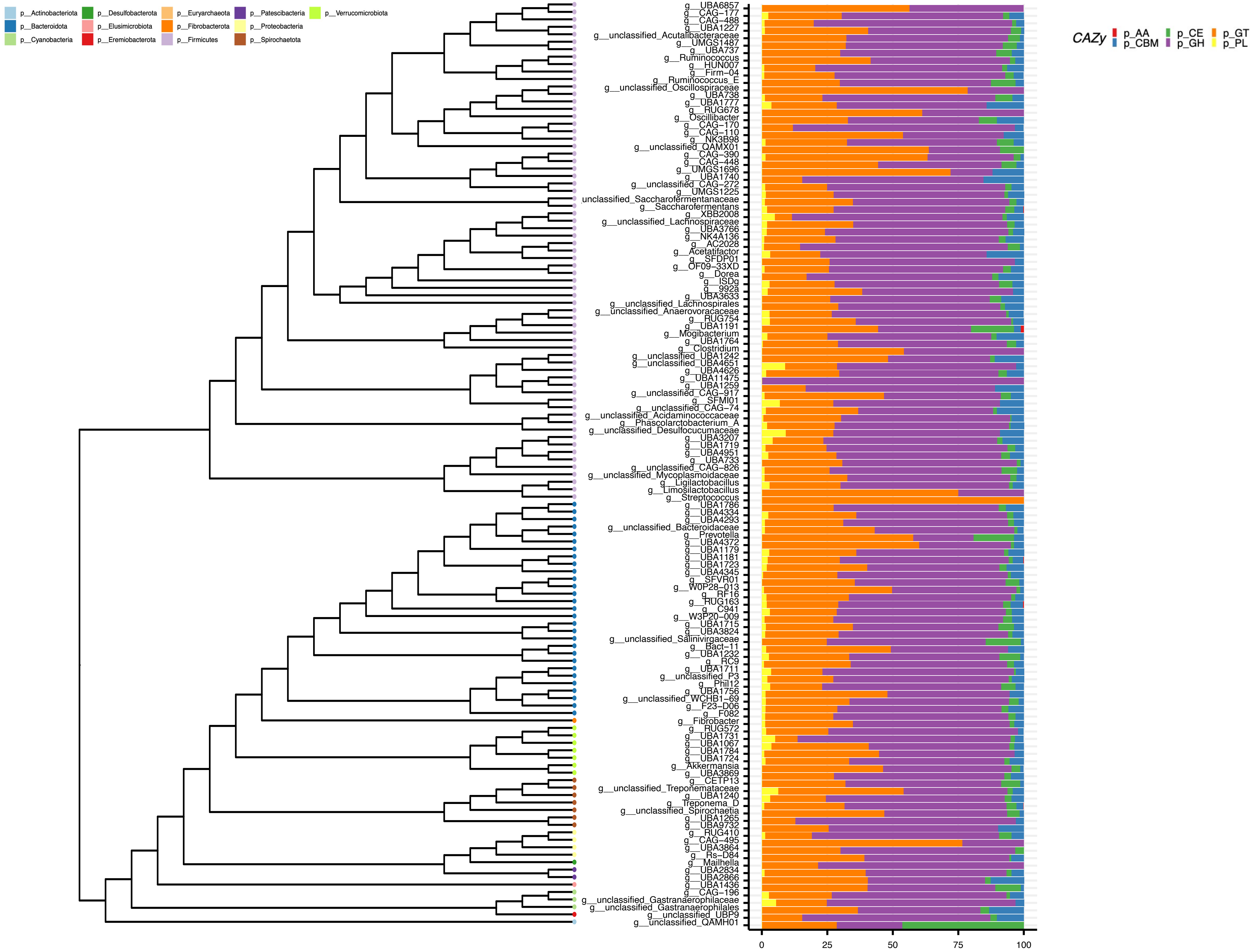

### Suppl Fig S11

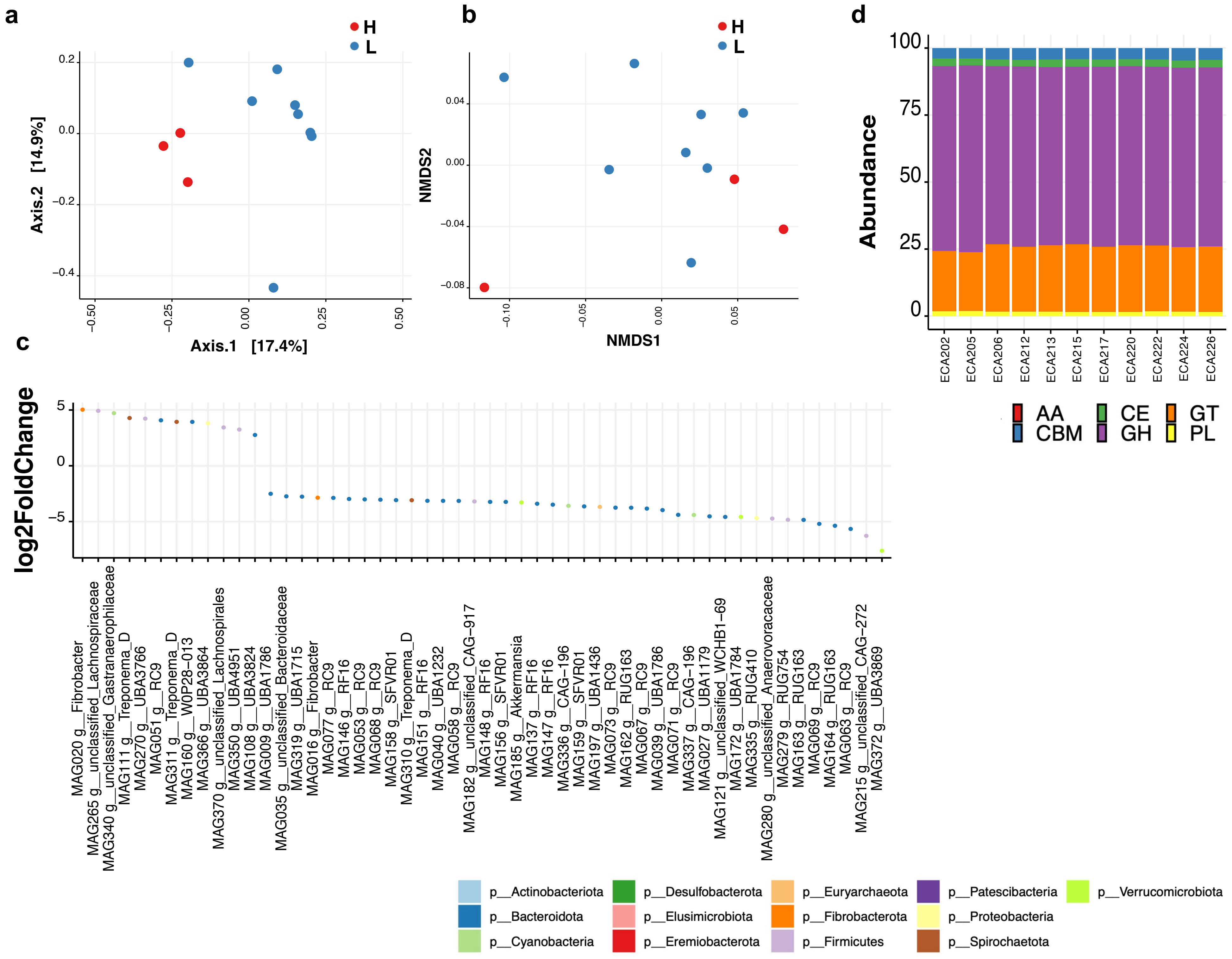
